# Inferring brain structural complexity from dynamical measures

**DOI:** 10.64898/2026.10.21.760743

**Authors:** Gianluca Gaglioti, Anna Cattani, Thierry Nieus, Davide Momi, Renzo Comolatti, Leonardo Dalla Porta, Simone Sarasso, Marcello Massimini

## Abstract

Complexity, commonly defined as the coexistence of integration and differentiation, is a fundamental property of thalamocortical networks and is regarded as a prerequisite for adaptive brain function and consciousness. A variety of metrics have been developed to estimate neural complexity from recorded activity and are increasingly employed in both research and clinical settings. Yet, the extent to which these measures reflect specific architectural features of the underlying circuits remains poorly understood. Here, we used Wilson-Cowan mean-field equations to simulate neural dynamics across networks spanning a broad range of architectures. By systematically rewiring connectivity, we generated graded variations in the structural integration–differentiation balance and assessed a panel of complexity measures derived either from spontaneous activity (observational metrics) or from responses to external perturbations (perturbational metrics). Observational measures exhibited markedly heterogeneous behavior: some were primarily sensitive to network integration, others to differentiation, whereas some showed limited sensitivity to architectural manipulations. In contrast, perturbational measures robustly peaked at intermediate rewiring steps and scaled with small-worldness, consistent with architectures that combine local recurrence and distributed communication. Moreover, perturbational metrics proved robust to common-driver confounds and were highest in a wake-like regime near critical instability, declining as sleep-like dynamics became more widespread across the network. Taken together, these findings map widely used empirical complexity measures onto specific network architectures, reveal dynamical factors that can bias observational estimates, and provide guidance for inferring complexity in systems whose internal organization is inaccessible, unconventional, or only partially known, including brain organoids, non-human nervous systems, and synthetic neural networks.

## 1. Introduction

A central question in neuroscience is how the brain’s extraordinarily rich repertoire of dynamics emerges from its underlying structural architecture (1). These dynamics are neither random nor trivially regular but instead express a hallmark property often referred to as complexity (2, 3). In this context, brain complexity is commonly framed as the coexistence of integration—reciprocal interactions across distributed regions—and differentiation—the preservation of locally specialized activity, structurally supported by network segregation (4, 5). Complexity is thought to be maximized when these two tendencies are optimally balanced. If integration dominates, activity becomes redundant and stereotypical; if differentiation dominates, activity fragments into weakly interacting modules. This balance may reflect an evolutionary cost-benefit optimum that supports adaptive flexibility by balancing efficient global integration with specialized local differentiation under strong constraints on wiring cost, space, and metabolism (6). Architectures that jointly maximize integration and differentiation have also been proposed as the optimal physical substrate for the unity and richness of conscious experience (7).

Graph-theoretic work and simplified models have identified several architectural motifs that can support this balance (8–12). At the structural level, integration is typically associated with distributed connectivity and high global efficiency, whereas differentiation is supported by network segregation, reflected in high local clustering. Small-world organization combines high clustering with short path lengths, enabling both local specialization and global communication (13, 14). Modularity and hierarchy promote specialization across subnetworks and interactions across multiple scales (15–18). Hubs and rich-club provide integrative convergence points for distributed processing (19–23), whereas spatial and metabolic constraints favor shorter links and local segregation (24, 25). At finer scales, recurrent connections and feedback loops create motifs that can sustain activity and support reverberant interactions, providing a structural basis for the emergence of diverse and metastable patterns of collective activity (26–30).

Although a balanced architecture combining local recurrence and distributed communication may provide the structural substrate for complexity, the emergence of complex dynamics also depends on the network’s operating regime (31, 32). From the perspective of dynamical systems and statistical physics, complexity has been proposed to be associated with regimes near the boundary between order and disorder (33–36). In these regimes, metastable dynamics enable flexible transitions between integrated and differentiated states (37, 38), while simultaneously supporting the computational advantages associated with near-critical dynamics (39–42). Importantly, network topology plays an active role in shaping these dynamical landscapes, influencing the location, breadth, and stability of near-critical regimes and thereby constraining the conditions under which complex dynamics can emerge (26, 28, 43–46).

Empirical neuroscience has developed a diverse toolkit of measures to estimate the integration-differentiation balance from brain activity, especially in humans (47, 48). These measures generally yield high values during wakefulness and other conscious conditions, such as dreaming and psychedelic or hallucinatory states, and lower values during states of diminished or absent consciousness such as sleep, anesthesia, and disorders of consciousness (49–61). Beyond global consciousness states, the integration-differentiation balance has also been linked to healthy cognition and to its disruption in aging and brain disorders (5, 62–70). Thus, complexity metrics are increasingly used not only as empirical descriptors, but also as practical markers of brain state and function.

Yet, despite their widespread use, the interpretation of these metrics remains uncertain because their link to specific underlying architectural motifs has not been systematically established. Structure–complexity relationships have been examined before, but mostly for single theoretical measures in abstract or linear models (2, 71); the empirically used panel has not been evaluated across a controlled structural sweep with the dynamical working point held fixed. A central distinction is between observational measures, which summarize ongoing activity and may reflect both intrinsic interactions and shared external drives, and perturbational measures, which quantify the spatiotemporal response to a controlled input. These distinct measurement logics suggest that the two classes may differ in their sensitivity to structural architecture, dynamical regime, and common-input confounds. Within each class, existing measures further rest on different assumptions and observables, ranging from entropy- and compressibility-based approaches to interaction-based metrics (48, 47). Consequently, it is often unclear whether different metrics capture the same underlying property or are preferentially sensitive to integration, differentiation, their optimal balance, near-critical operating regimes, or even confounding factors such as common input. Resolving these ambiguities is essential for interpreting empirical results and for applying complexity measures to systems whose internal organization is unknown.

Here we use computational modeling to bridge graph theory and empirical complexity metrics in a controlled, mechanistic framework. We employ a mean-field model of coupled excitatory and inhibitory populations and simulate dynamics on networks whose architecture is systematically rewired to span graded regimes of structural integration and segregation. We then apply both observational and perturbational complexity measures to determine how each responds to specific structural modifications. We further challenge these measures with two distinct dynamical manipulations: a progressive transition from wake-like toward globally synchronous sleep-like dynamics, representing a genuine alteration of the network’s operating regime, and the introduction of a common driver, serving as a controlled dynamical confound. By explicitly linking network motifs to complexity estimates, this approach clarifies what current metrics can (and cannot) infer about the underlying architecture. Also, it provides a principled basis for interpreting complexity in systems where the structure is less accessible or less canonical—such as organoids, non-human brains, and synthetic or engineered neural networks.

## 2. Results

### 2.1 Setting the dynamical working point across topologies

We generated two complementary network families traversing the continuum between segregation and integration: a ring-to-random family (C1 → B1 → R; randomization) and a random-to-modular family (R → B2 → C2; modularization), both traversing an intermediate small-world regime characterized by the coexistence of high clustering and high efficiency (Fig. 1A-B).

**Figure 1.**
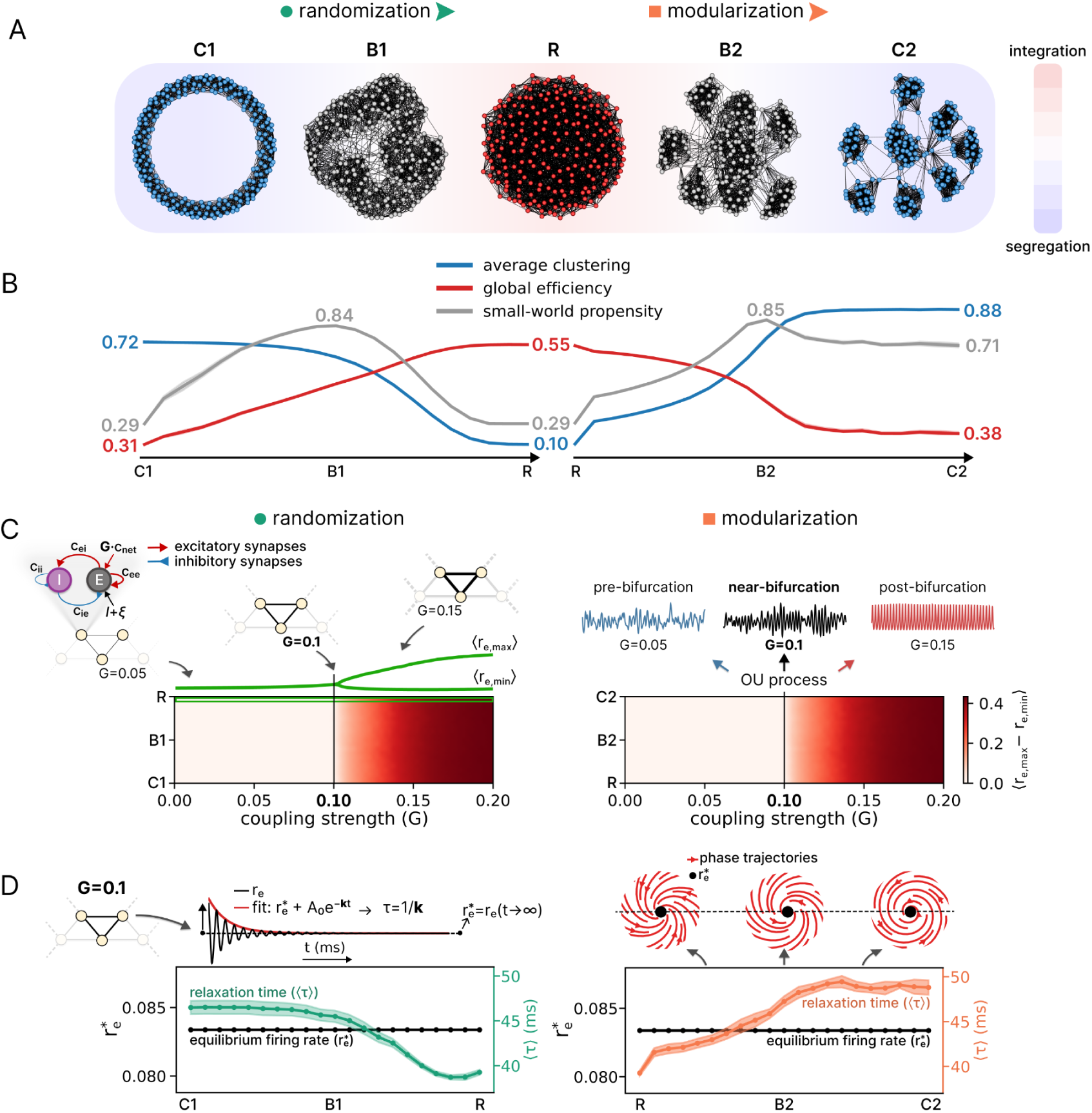
Network topologies and dynamics. **(A)** Starting from a regular ring lattice (C1, top left), connections were randomized, yielding small-world topologies (B1) and eventually a random network (R, top right). From the random network, a modularization algorithm was applied yielding balanced modular networks (B2) and finally a strongly modular network (C2, bottom right). The rewiring procedure spans architectures of predominant integration (i), segregation (s), or i/s balance. **(B)** Network measures across rewiring steps. Randomization (top panel) decreased average clustering (blue line) and increased global efficiency (red line), with a peak in small-world propensity (purple line) at intermediate steps (B1). Modularization (bottom panel) showed the complementary trend, again passing through a balanced regime (B2). **(C)** Each network node was modeled using Wilson–Cowan equations (inset). Deterministic simulations were performed while varying the global coupling strength G from 0 to 0.2 in increments of 0.0025. Heat maps show the node-averaged difference between the maximum and minimum excitatory firing rates (⟨ r_e,max_ − r_e,min_⟩), as a function of G. This quantity is zero when the dynamics converge to a fixed point and becomes nonzero when sustained oscillations emerge. Across topologies, it increased just above G≈0.1, where a supercritical Andronov–Hopf bifurcation gave rise to a stable limit cycle. The green trace above the left heat map show r_e,max_ and r_e,min_ (averaged across nodes ⟨·⟩) for the random network. The right panels show representative trajectories after addition of an Ornstein–Uhlenbeck process: G=0.05, pre-bifurcation noisy fluctuations (blue); G=0.1, near-bifurcation noise-driven damped gamma oscillations (black); and G=0.15, post-bifurcation sustained oscillations (red). (**D)** Relaxation-time estimation at G=0.1. For each network, a small displacement from the equilibrium excitatory firing rate, r_e_∗, was applied to each sampled node. The successive positive peaks of the return trajectory were fitted with an offset exponential decay, r_e_∗+A_0_e^−kt^ (red), and the relaxation time was defined as τ=1/k. This procedure was repeated for 10 nodes per network, and the node-averaged relaxation time, ⟨τ⟩, was used for subsequent analyses. The upper-right panels show schematic phase portraits illustrating topology-dependent differences in the return trajectories toward the fixed point (red curves).

To simulate neuronal network dynamics, each node was modeled according to the Wilson-Cowan equations, simulating the interaction between excitatory and inhibitory populations (Fig. 1C). The deterministic system was characterized by varying the global coupling strength G (from 0 to 0.2 in increments of 0.0025), encompassing a quiescent regime followed by a bifurcation giving rise to stable oscillations. Because rewiring preserved the in-degree of every node, changing G scaled the total recurrent input to each node equivalently across topologies. Accordingly, in the quiescent regime the networks shared the same equilibrium firing rate (r_e_*) for each G, so differences in their dynamics could be attributed to topology rather than to differences in operating point. Across topologies, increasing G produced a similar qualitative bifurcation structure: the equilibrium fixed point lost stability just above G≈0.1, giving rise to a stable limit cycle through a supercritical Andronov–Hopf bifurcation. We therefore set G=0.1, just below this transition, for subsequent simulations. At this operating point, the addition of Ornstein–Uhlenbeck fluctuations produced noise-driven damped gamma oscillations (Fig. 1C, right panel, black trace) and maximized coordinated network fluctuations (Supplementary Fig. S2), consistent with metastable dynamics near critical instabilities (36).

Although identical in-degree guarantees a common r_e_*, the underlying connectivity structure can differentially modulate the local geometry of the attractor. To quantify this topological influence on dynamical stability, we estimated the effective relaxation time (τ) of small deviations from r_e_* using a peak-fitting procedure applied to the damped oscillations (Fig. 1D). Across the segregation–integration continuum, the mean τ (⟨τ⟩) varied systematically with topology. Networks with high clustering exhibited markedly longer ⟨τ⟩, whereas random networks recovered more rapidly (Fig. 1D). This pattern indicates that clustered motifs act as reverberating circuits in which activity circulates through recurrent loops (29), sustaining perturbations and drawing the system closer to its stability boundary. Conversely, the absence of such systematic loop amplification in random topologies promotes faster activity dissipation, conferring a greater stability margin (Fig 1D, right panel, red trajectories). These findings confirm that for a given equilibrium firing rate, topology shapes the attractor landscape, with recurrent connectivity slowing down dynamics and shifting networks closer to critical instability.

### 2.2 Spontaneous and evoked dynamics reflect structural topological constraints

We next characterized spontaneous interactions by analyzing functional connectivity (FC). Representative FC matrices (Fig. 2A) showed that clustered networks closely reflected their underlying structural organization: the ring lattice (C1) displayed strong nearest-neighbor correlations, whereas the modular network (C2) showed strong within-module correlations. Balanced networks (B1, B2) preserved these clustering signatures, while the random network (R) exhibited a diffuse, diluted FC matrix with no spatial structure. These patterns were confirmed by the FC distributions, which were long-tailed in clustered and balanced networks but narrower in the random network (Fig. 2B). Across rewiring (Supplementary Fig. S3A), randomization progressively reduced the upper tail of the FC distribution while raising its median, indicating more spatially diffuse interactions around a higher baseline; modularization produced the opposite effect, with strong within-cluster correlations (higher 95th percentile) and weak between-cluster correlations (lower 5th percentile). Consistent with these trends, the median FC correlated strongly with global efficiency (Fig. 2C; randomization: ρ=0.86; modularization: ρ=0.87), whereas the 95th percentile correlated with average clustering (Fig. 2D; randomization: ρ=0.86; modularization: ρ=0.87), confirming that clustered motifs reinforce local pairwise interactions, whereas random architectures promote widespread but diluted coupling.

**Figure 2.**
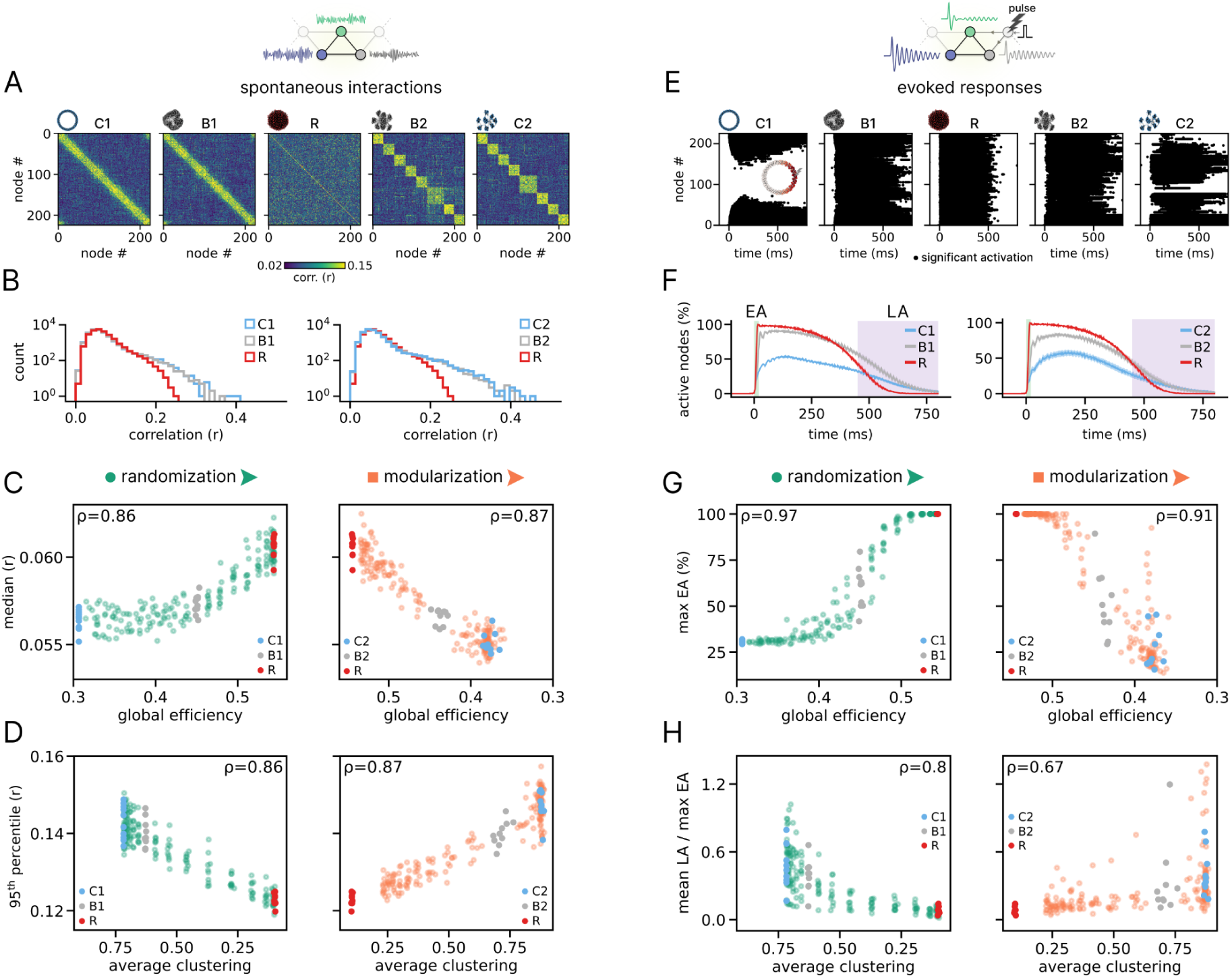
Spontaneous and evoked activity signatures across network families. **(A)** Functional connectivity (FC) matrices were computed from spontaneous activity in representative networks (C1, B1, R, B2, C2) along the randomization and modularization processes. FC was estimated as the maximum cross-correlation across lags for each node pair. For modular networks (B2, C2), node indices were resorted by community membership, identified via the Louvain algorithm on the structural graph. Example traces illustrate spontaneous fluctuations in excitatory population activity (r_E_). **(B)** Distribution of FC correlation values for representative networks. Clustered and balanced networks exhibit a pronounced right tail of high correlations, absent in the random network. **(C)** Median FC correlation as a function of global efficiency across rewiring steps, for both network families (randomization: ρ=0.86, p=2×10^-63^; modularization: ρ=0.87, p=6×10^-65^). **(D)** 95th percentile of the FC distribution as a function of average clustering (randomization: ρ=0.86, p=2×10^-61^; modularization: ρ=0.87, p=5×10^-66^). **(E)** Spatiotemporal matrices of significant node activations following an excitatory pulse delivered to a single node (inset schematic), shown for representative networks along both rewiring processes. Superimposed traces illustrate trial-averaged evoked responses (r_E_) for the three nodes with the largest early activation (assessed by the area under the response envelope in the first 20 ms). **(F)** Time courses of the percentage of significantly active nodes (mean ± SEM across network realizations). EA and LA denote the early (0-20 ms; green shaded area) and late (450-800 ms, purple shaded area) activation windows, respectively. **(G)** Maximum early activation (max_EA_) as a function of global efficiency (randomization: ρ=0.97, p=6×10^-126^; modularization: ρ=0.91, p=5×10^-80^). **(H)** Ratio of mean late to maximum early activation (mean_LA_/max_EA_) as a function of average clustering (randomization: ρ=0.80, p=2×10^-47^; modularization: ρ=0.67, p=2×10^-28^). In all scatter plots, each point represents a single network realization (10 per rewiring step); colors identify the representative networks (C1, B1, R, B2, C2). Shaded areas denote SEM across network realizations.

We then asked whether perturbation would better reveal the balanced regime, which should combine integration and differentiation in the evoked responses (Fig. 2E). Following a brief stimulation of a single node, clustered architectures (C1, C2) produced sustained but spatially confined responses, while the random network (R) produced network-wide but stereotypical and short-lived responses. Balanced architectures combined network-wide and sustained responses, consistent with the coexistence of clustered recurrent motifs supporting temporal amplification and network shortcuts supporting widespread propagation (Fig. 2F). Accordingly, maximum early activation (max_EA_) decreased from random to clustered networks, mean late activation (mean_LA_) peaked in balanced configurations, and the mean_LA_/max_EA_ ratio increased in clustered architectures as clustering promotes temporal reverberation at the expense of the initial spatial propagation (Supplementary Fig. S3B). Consistently, these metrics were strongly predicted by network topology (Fig. 2G–H): max_EA_ correlated with global efficiency (randomization: ρ=0.97; modularization: ρ=0.91), while mean_LA_/max_EA_ correlated with average clustering (randomization: ρ=0.80; modularization: ρ=0.67).

### 2.3 Dynamical measures of complexity reveal different sensitivities to network segregation and integration

We next quantified the complexity of both spontaneous and evoked activity using observational and perturbational metrics, respectively (Supplementary Fig. S1). For spontaneous activity, we assessed complexity using two classes of measures: signal diversity and interaction-based complexity. Signal diversity was quantified using Lempel-Ziv complexity (LZc), amplitude coalition entropy (ACE), and synchrony coalition entropy (SCE). Interaction-based complexity was characterized by neural complexity (NC) and functional complexity (F_comp_). For evoked activity, we computed the perturbational complexity index (PCI), the evoked pattern dimensionality, the spatiotemporal entropy (i.e., spatial, temporal, and their average ⟨H⟩), and the entropy rate averaged over nodes (H_rate_). The evolution of these measures across rewiring steps is shown in Fig. 3.

**Figure 3.**
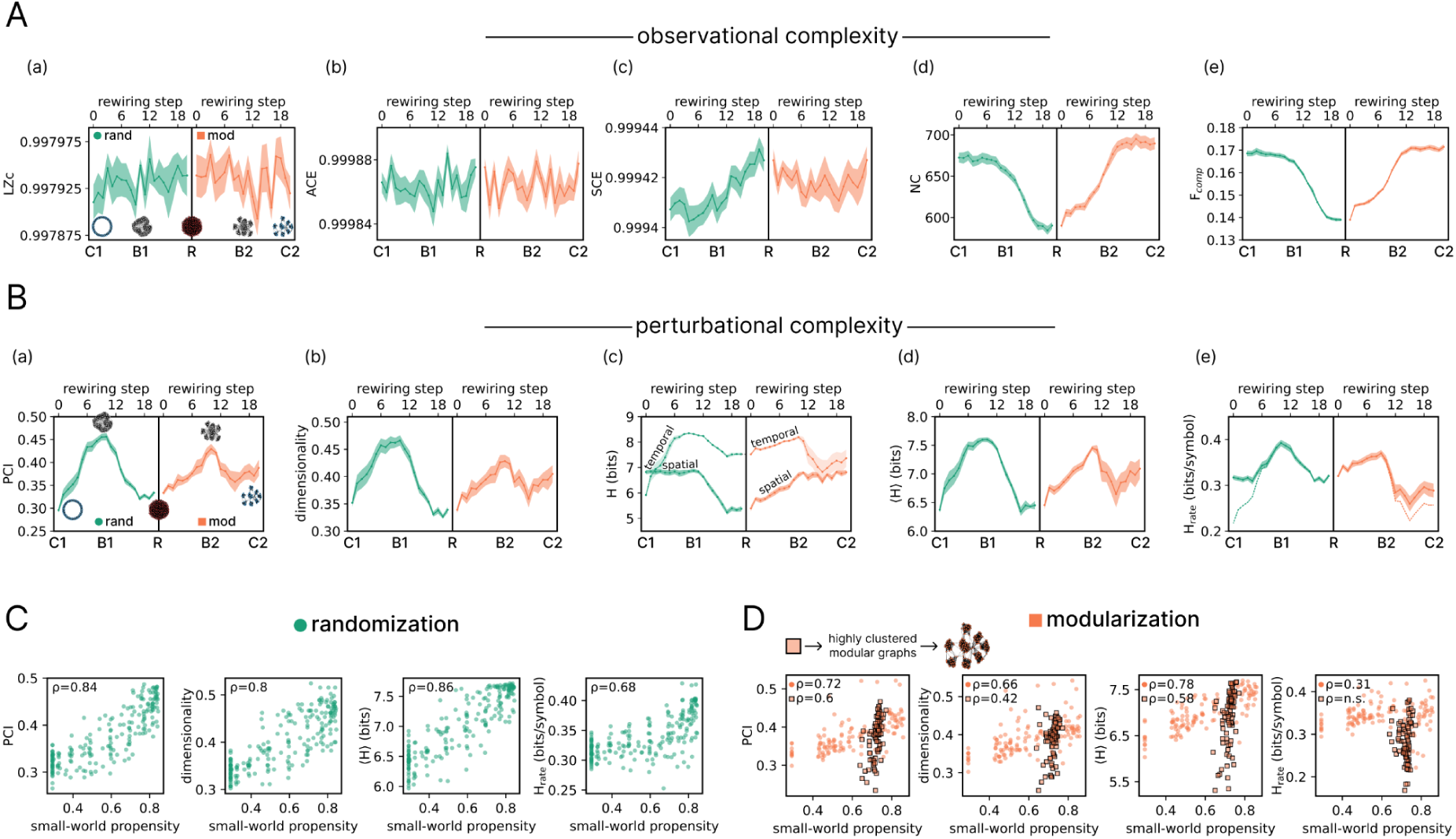
Different sensitivities of dynamical measures to structural complexity. **(A)** Observational complexity metrics computed from spontaneous activity, shown as a function of rewiring step for both the randomization (green) and modularization (orange) processes: Lempel-Ziv complexity (LZc; panel a), amplitude coalition entropy (ACE; b), synchrony coalition entropy (SCE; c), neural complexity (NC; d), and functional complexity (F_comp_; e). **(B)** Perturbational complexity metrics computed from evoked responses: perturbational complexity index (PCI; panel a), pattern dimensionality (dimensionality; b), entropy of spatial and temporal patterns (H; c), their average (⟨H⟩; d), and entropy rate (H_rate_; e). In panel e, dashed lines indicate the average entropy rate computed including nodes with no significant evoked response, which contribute zero entropy rate. In all panels, lines and shaded areas represent the mean ± SEM across network realizations (n=10 per step); network topology icons along the x-axis identify representative networks at selected rewiring steps (C1, B1, R for the randomization process; R, B2, C2 for the modularization process). **(C)** Scatter plots showing perturbational complexity measures (PCI, dimensionality, ⟨H⟩, and H_rate_) as a function of small-world propensity across rewiring steps of the randomization process. Spearman correlation coefficients (ρ) are reported for each metric. **(D)** Same as (C) for the modularization process. Highly clustered modular networks (identified by a threshold on clustering coefficient values; see Supplementary Fig. S8) are highlighted as square markers. Correlation coefficients are reported separately for the highly clustered subgroup (squares) and the rest of the networks (circles); n.s. indicates non-significant correlation (p>0.05).

Overall, observational metrics exhibited weaker and more heterogeneous sensitivity to network architecture than perturbational measures. LZc and ACE did not vary significantly across rewiring steps in either process (repeated-measures ANOVA, all p > 0.05), although LZc showed a slight tendency to increase with greater network integration. This tendency became more apparent in the supplementary analyses across different network sizes and densities, where similar trends were also visible for ACE and SCE (Supplementary Fig. S4C). In the main analysis, SCE increased along the randomization process but reached a plateau from R to C2 during modularization (Fig. 3A, c). NC and F_comp_ showed similar profiles, both increasing toward more clustered architectures, consistent with their sensitivity to the structure of network interactions (Fig. 3A, d–e). In contrast, all perturbational metrics followed a pronounced inverted-U profile across both rewiring processes, peaking around B1 and B2, where integration and segregation were balanced (Fig. 3B, a–e), with the same pattern preserved across different network sizes and densities, stimulation amplitudes, and delay distributions (Supplementary Figs. S4D, S5, S6).

Next, we assessed the interrelationships among structural, functional, and complexity dynamical measures by computing their Spearman correlations across all networks (Supplementary Fig. S7). These analyses confirmed a robust correlation between perturbational complexity metrics and the small-world propensity of the underlying graphs (Fig. 3C-D). During the modularization process (Fig. 3D), we identified a subset of highly clustered networks in which perturbational complexity remained markedly variable despite very similar global topology. We isolated this subgroup using a threshold on the clustering coefficient (Supplementary Fig. S8). Within this restricted set, complexity remained significantly related to small-world propensity, with the exception of entropy rate. Global efficiency showed little explanatory power, whereas local neighborhood efficiency of the stimulated node better accounted for variability in perturbational complexity (Supplementary Fig. S9), highlighting the role of connector-hub-like neighborhoods in shaping complex responses.

Because topology also influenced relaxation dynamics (Fig. 1D), we tested whether the perturbational complexity peak could be explained by differences in proximity to the stability boundary (Supplementary Note 4.1; Supplementary Figs. S10, S11). PCI remained higher in balanced than in random networks before differences in decay could dominate (Supplementary Fig. S11A–B), and perturbational complexity remained positively associated with small-world propensity after matching networks for relaxation time (Supplementary Fig. S11C–D). Thus, attractor stability contributes to, but does not fully explain, the topology–complexity relationship (see Supplementary Note 4.2).

### 2.5 Divergent sensitivities of complexity metrics to common driver confounds

To assess the robustness of complexity metrics to a major confound in empirical data, i.e., shared input, we introduced a common external driver to all nodes and systematically increased its strength while keeping total input variance constant (Fig. 4A). Shared inputs, such as neuromodulatory fluctuations or sensory stimulation, can induce widespread coordination across structurally independent nodes, artificially inflating functional integration and reducing functional differentiation. We evaluated the effects of this manipulation across representative network classes (C1, B1, R, B2, C2). As expected, the median value of the FC distribution monotonically increased with common driver strength *s* (Fig. 4B), confirming the emergence of spurious pairwise correlations independent of network topology.

**Figure 4.**
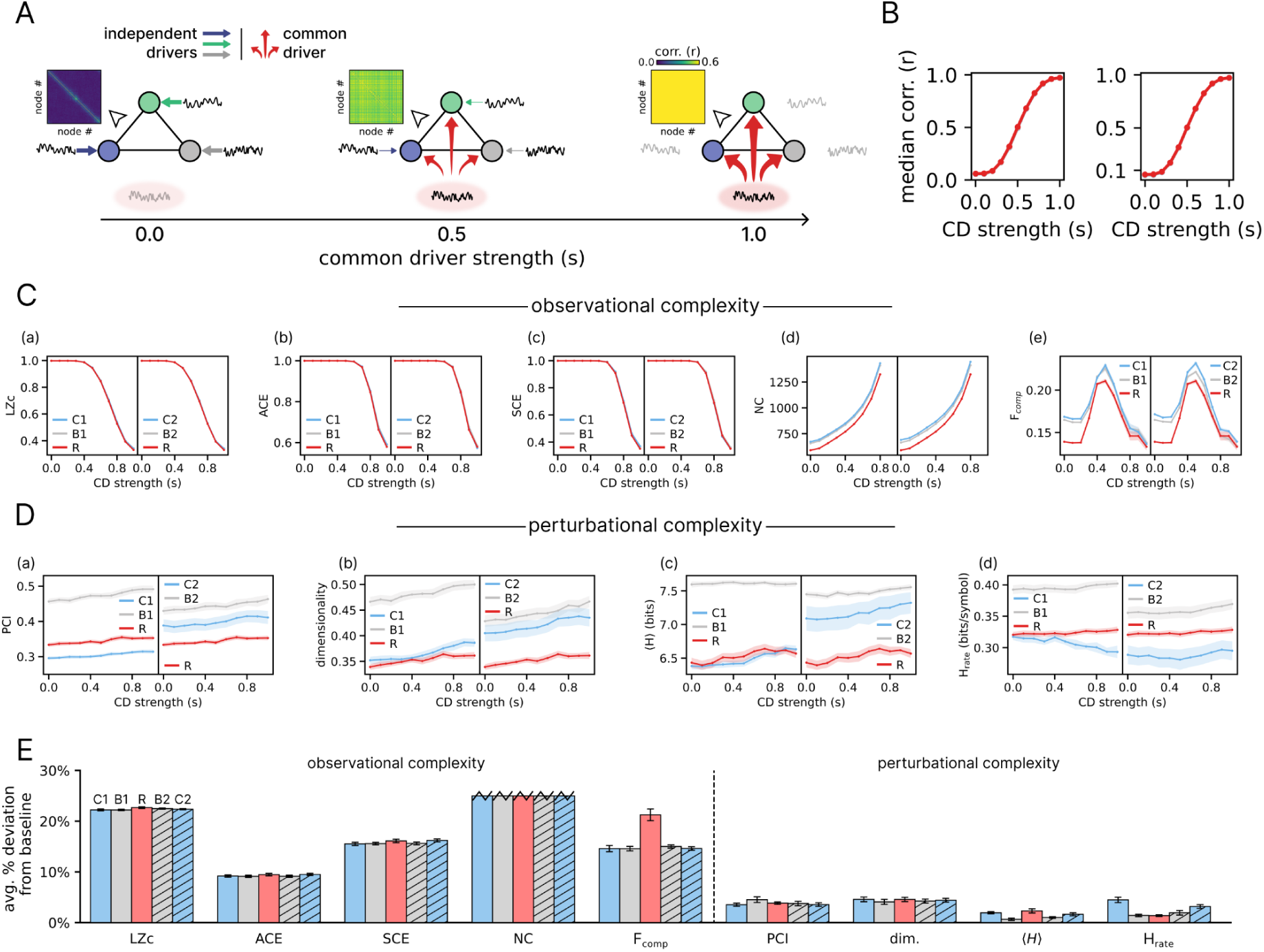
Different sensitivities of dynamical measures of complexity to common driver confounds. **(A)** Schematic of the common driver protocol. Each node received an independent Ornstein-Uhlenbeck (OU) input, and a shared OU process was added with mixing strength *s*, ranging from 0 (independent inputs only) to 1 (common input only). Representative examples for increasing *s* are shown with the corresponding functional connectivity (FC) matrices, illustrating the increase in pairwise correlations induced by the common driver. **(B)** Median FC correlation as a function of common driver strength *s* for the representative network classes of the randomization-family (left: C1, B1, R) and the modularization-family (right: R, B2, C2). In both families, FC correlation increases monotonically with s. **(C)** Observational complexity metrics computed from spontaneous activity as a function of *s*: Lempel-Ziv complexity (LZc; a), amplitude coalition entropy (ACE; b), synchrony coalition entropy (SCE; c), neural complexity (NC; d), and functional complexity (F_comp_; e). Left and right subpanels show the randomization-family (C1, B1, R) and modularization-family (C2, B2, R) network classes, respectively. NC values beyond *s=*0.8 are omitted because estimates became numerically unreliable due to singular or near-singular correlation matrices. **(D)** Perturbational complexity metrics computed from evoked responses as a function of *s*: perturbational complexity index (PCI; a), pattern dimensionality (dimensionality; b), mean entropy of the spatial and temporal patterns (⟨H⟩; c), and entropy rate averaged across nodes (H_rate_; d). Left and right subpanels show the randomization-family (C1, B1, R) and modularization-family (C2, B2, R) network classes, respectively. Curves represent the mean across network realizations (n=10); shaded areas indicate SEM. For H_rate_, nodes with no significant evoked response, which contribute zero entropy rate, were excluded from the average. **(E)** Common driver sensitivity for each metric, quantified as the mean absolute percentage deviation from the *s=*0 baseline across all common driver strengths. Bars show mean ± SEM across network realizations (n=10) per network class. NC is capped at 25% for visualization.

We then assessed the deviation of each complexity metric from its baseline value (*s*=0). Spontaneous activity metrics proved highly vulnerable, although with heterogeneous directions of sensitivity. Measures of signal diversity, including LZc, ACE, and SCE, progressively decreased as spurious connectivity increased (Fig. 4C, a-c). In contrast, NC systematically increased with *s* (Fig. 4C, d), reflecting its bias toward integration. F_comp_ exhibited a non-monotonic profile, peaking at intermediate values of *s*, where independent and shared fluctuations were balanced (Fig. 4C, e), consistent with its dependence on the integration–differentiation balance but also highlighting its sensitivity to spurious activity sources.

By contrast, perturbational complexity exhibited greater robustness. Given their causal nature, these metrics better preserved their ability to discriminate between network classes despite increasing common driver strength (Fig. 4D, a-d). Moreover, across all tested values of *s*, perturbational metrics consistently yielded higher separation between network classes. To quantify and compare the sensitivity to common drivers, we computed for each metric the mean absolute percentage deviation from the *s*=0 baseline across all common driver strengths (Fig. 4E). Observational metrics showed substantially higher sensitivity (LZc: 22.4%; ACE: 9.3%; SCE: 15.8%; NC: 43.0%; F_comp_: 16.0%) than perturbational ones, with the latter all remaining below 5% average deviation across network classes (PCI: 3.8%; dimensionality: 4.4%; ⟨H⟩: 1.6%; H_rate_: 2.5%).

### 2.5 Sleep-like dynamical transitions modulate network complexity

The brain can transition between profoundly different functional states, such as wakefulness and NREM sleep, while its structural connectivity remains largely unchanged. We therefore asked whether the complexity supported by a given network architecture also depends on the dynamical regime in which that architecture operates. We focused on the brain-like balanced modular networks (B2) and used an extension of the Wilson–Cowan model incorporating spike-frequency adaptation (72, 73), which can reproduce transitions from wake-like to sleep-like activity. We adopted *g*=1.8 as the baseline condition (as in 73) and doubled adaptation to *g*=3.6 to induce sleep-like dynamics. We first sampled a node and identified its community as the reference module. Starting from this module, sleep-like dynamics were progressively imposed on additional modules, producing a graded transition from locally restricted to network-wide sleep-like activity while leaving structural connectivity unchanged (Fig. 5A).

**Figure 5.**
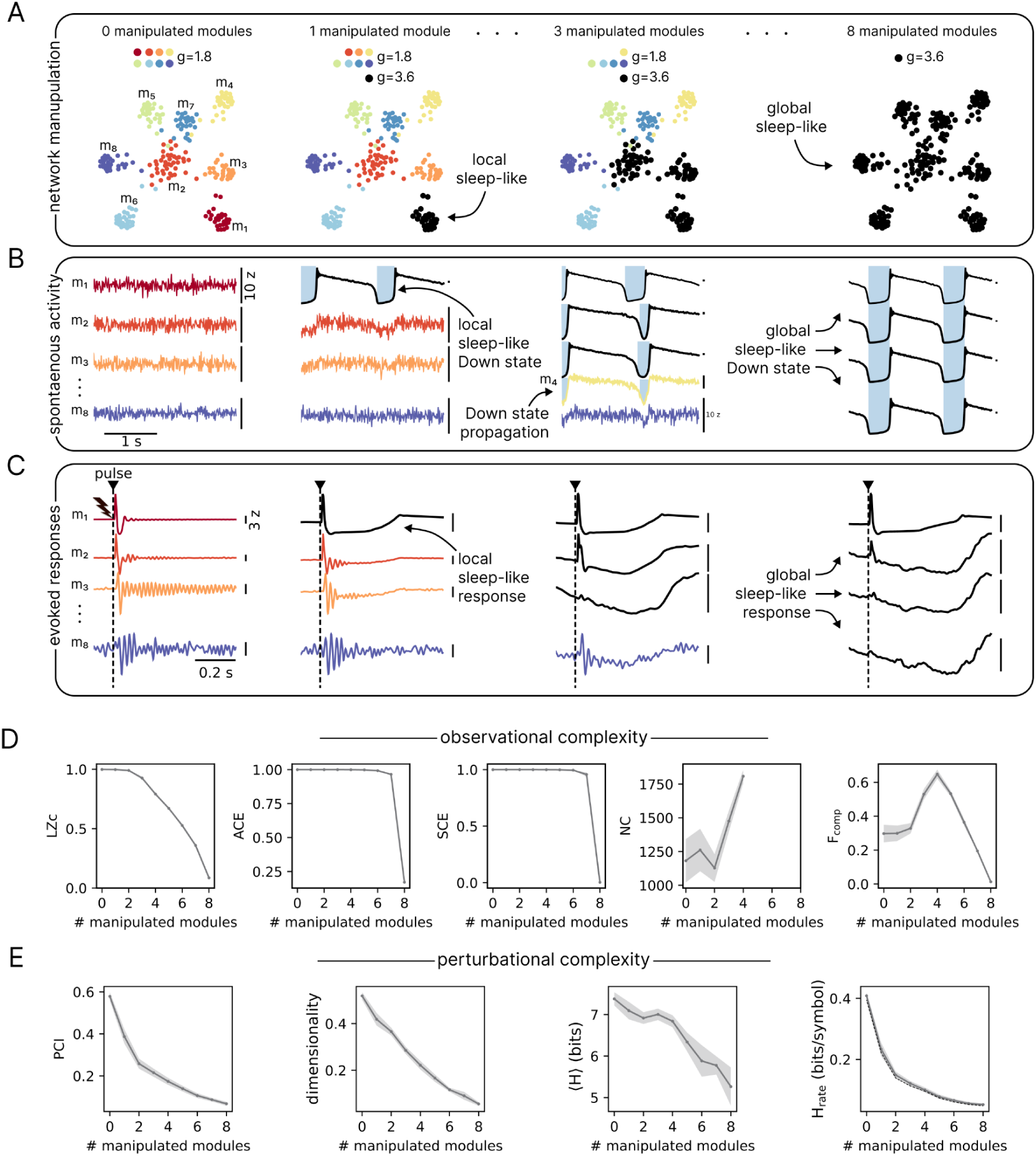
Dynamical regime modulates network complexity. **(A)** Progressive transition from wake-like *g=*1.8 to sleep-like *g=*3.6 dynamics in a representative B2 network. A sampled node defined the reference module; subsequent modules were manipulated in descending order of their mean structural connectivity with the reference module. Black nodes indicate manipulated modules. **(B)** Representative spontaneous activity, averaged across nodes within each displayed module. Down states emerge locally in manipulated modules; with three modules manipulated, a Down state deflection also appears in non-manipulated m_4_, consistent with propagation. Down states become global when all modules are manipulated. Traces were z-scored relative to the unmanipulated condition to preserve amplitude differences across manipulations. The vertical scale bar on the right corresponds to 10 z-score units. **(C)** Representative module-averaged evoked responses following stimulation of the sampled node. Fast distributed reverberations progressively disappear as sleep-like dynamics spread across the network. Each trace was independently z-scored to emphasize response temporal dynamics rather than absolute amplitude. The vertical scale bar corresponds to 3 z-score units. **(D)** Observational complexity metrics across manipulation steps. NC values beyond four manipulated modules are omitted because estimates became numerically unreliable due to singular or near-singular correlation matrices. **(E)** Perturbational complexity metrics progressively decrease as sleep-like dynamics become more widespread. For H_rate_, the dashed black line includes nodes with no significant evoked response as zero-valued contributions. Lines and shaded areas indicate the mean ± SEM across network realizations. Seven of the ten B2 networks contained eight modules, whereas three contained nine; analyses were therefore restricted to the seven eight-module networks to ensure comparison across the same number of manipulation steps.

This manipulation progressively transformed spontaneous activity (Fig. 5B). The fully wake-like network displayed irregular, low-amplitude fluctuations around a fixed point near oscillatory instability. When only the reference module was shifted to high adaptation, local Down states emerged while the remaining modules largely retained wake-like activity, with some Down states deflections consistent with propagation from the manipulated module. As additional modules entered the sleep-like regime, slow fluctuations became increasingly widespread and coordinated, culminating in global Down states.

We next stimulated the sampled node (Fig. 5C). In the wake-like regime, stimulation elicited rich, long-lasting reverberations comprising multiple fast response components across modules. Once the stimulated module entered the sleep-like regime, its response became dominated by a slower component and complex reverberation was markedly reduced. As sleep-like dynamics extended across the network, fast response components were progressively lost also in modules that had not yet been manipulated. When all modules were in the sleep-like regime, stimulation evoked predominantly network slow, stereotyped responses.

Observational complexity measures showed heterogeneous sensitivity to the manipulation (Fig. 5D). Signal-diversity measures generally decreased: LZc declined progressively, whereas ACE and SCE remained relatively preserved initially before decreasing at later stages. In contrast, NC increased as activity became increasingly synchronous with progressive manipulation, whereas F_comp_ peaked at intermediate stages, when wake-like and sleep-like modules coexisted and the FC distribution was broadest.

Perturbational complexity showed a more consistent dependence on dynamical state (Fig. 5E), with all metrics decreasing as sleep-like dynamics became more widespread and evoked responses increasingly stereotyped. Thus, despite unchanged structural architecture, the network’s capacity to express a rich repertoire of causal interactions depended strongly on its dynamical regime and progressively declined as sleep-like dynamics extended from local to global scales.

## 3. Discussion

This study clarifies how different activity-based measures of complexity relate to the structural organization of neuronal networks across dynamical regimes. Using a controlled mean-field model, we show that complexity metrics are not interchangeable: some primarily reflect structural integration, others differentiation, while others are strongly affected by external confounds. A central implication is that empirical complexity metrics should not be treated as an homogeneous class, but rather as complementary measures that support different levels of inference about the underlying system.

Among observational complexity measures, the interaction-based metrics NC and F_comp_ both increased toward more clustered architectures (Fig. 3A). This pattern can be understood in terms of the changes in spontaneous functional connectivity (FC) induced by network rewiring (Fig. 2A–D). Random graphs exhibited widespread but relatively weak pairwise interactions, whereas clustered networks combined strong within-cluster coupling with weak between-cluster coupling, resulting in a broader and more heterogeneous FC distribution. For F_comp_, the increase reflected this greater heterogeneity of pairwise FC, while the parallel increase in NC suggests that clustering also enriched multivariate dependencies across network partitions.

Signal diversity metrics, including LZc, ACE, and SCE, showed comparatively little sensitivity to topological rewiring. These measures capture the richness of amplitude or synchrony patterns rather than the organization of interregional statistical dependencies. Indeed, all networks operated within a near-bifurcation regime, preserving rich endogenous fluctuations across architectures irrespective of topology. Thus, the observational metrics tested here captured distinct aspects of spontaneous dynamics without consistently identifying the architectures that balanced structural integration and segregation.

Perturbational measures were more consistently associated with this structural balance. When the system was actively perturbed, evoked responses varied systematically across topologies (Fig. 2E). In random networks, stimulation produced rapid network-wide recruitment but limited sustained reverberation. In clustered networks, responses remained spatially confined but temporally persistent, consistent with local amplification within densely connected clusters (26, 28, 29). Balanced architectures combined both properties: inter-community shortcuts enabled rapid spatial propagation, whereas intra-community recurrence sustained temporal amplification (Fig. 2E–H). Accordingly, perturbational complexity followed a robust inverted-U profile, peaking in intermediate, small-world-like networks (Fig. 3B). Part of this effect could reflect topology-dependent changes in dynamical regime (26, 28, 29, 43, 44, 46), as clustered architectures relaxed more slowly and lay closer to the stability boundary. However, this did not fully account for the complexity peak: after matching networks for relaxation time, perturbational complexity remained positively associated with small-world propensity (Supplementary Fig. S11).

These findings help bridge graph theory and empirical measures of brain complexity. Small-world architectures have long been proposed as structural solutions to the integration–differentiation trade-off (6). Here, perturbational complexity provides a dynamical expression of this structural principle, highlighting the importance of dense local recurrent connectivity alongside efficient long-range communication. The dense, recurrent, and patchy organization of local cortical connectivity supports differentiated amplification through local feedback loop (27, 74–77), while short paths enable rapid network-wide propagation (13, 78, 79). Our results show that these properties jointly maximize perturbational complexity, expanding the repertoire of spatiotemporal patterns that can be recruited by perturbation.

Architecture, however, does not uniquely determine the complexity expressed by a network. Empirical data suggest that complexity can change and collapse even when the underlying structural architecture remains unchanged, as observed in healthy human brains during transitions from wakefulness to sleep or under general anesthesia (50, 52, 53, 60, 80, 81). To examine this state dependence, we held the structurally optimal architecture fixed, while progressively increasing spike-frequency adaptation; this activity-dependent mechanism is known to play key role in shifting activity from asynchronous, wake-like dynamics toward sleep-like regimes dominated by slow waves and by the associated neuronal downstates (82–84). As progressively more modules entered the sleep-like regime, perturbational complexity declined (Fig. 5). In the wake-like regime, recurrent connections supported rich and prolonged reverberations; with increasing adaptation, Down states interrupted this feedback and progressively obliterated fast evoked components. The same anatomical connections could therefore support different repertoires of effective interactions depending on the responsiveness of the neuronal populations they linked. Structural connectivity constrains the possible interactions, whereas the dynamical regime determines which can be expressed.

This manipulation also revealed convergence between observational and perturbational measures. Signal diversity metrics also decreased as sleep-like dynamics became more widespread, consistent with a shared sensitivity to the restriction of the network’s dynamical repertoire. This agrees with empirical findings showing reduced spontaneous signal diversity during general anesthesia and NREM sleep compared with wakefulness (53, 81, 85). The different responses to topological rewiring and to adaptation illustrate why agreement between measures in one setting does not imply that they capture the same property in another.

A further distinction concerns sensitivity to influences that alter recorded activity without changing intrinsic network connectivity. To explore this issue we progressively increased the influence of a common external driver over intrinsic network interactions (Fig. 4). The common driver increased apparent functional coupling and altered observational complexity measures, whereas perturbational measures remained comparatively robust and continued to discriminate network architectures even when spontaneous dynamics were dominated by the external drive. In our simulations, averaging across repeated stimulation trials attenuated activity that was not consistently time-locked to the perturbation, allowing the evoked response to reveal intrinsic network interactions. This mechanism helps explain the robustness of perturbational measures, but does not imply immunity to all confounds: external influences that systematically affect evoked responses may survive averaging. Observational measures remain informative about the activity the system expresses, although that activity may reflect both intrinsic interactions and external influences, including common input, sensory drive, volume conduction, or extracerebral sources (86–88).

These distinctions offer a framework for interpreting empirical dissociations between spontaneous and perturbational complexity. In minimally conscious patients, for example, severely abnormal spontaneous EEG can coexist with high perturbational complexity (89). Slow activity from local sources may dominate scalp recordings through volume conduction, while stimulation still recruits rich interactions within preserved circuits. Similarly, neuromuscular blockade reveals a strong effect of muscle activity on spontaneous metrics during anesthesia (90) whereas averaging can reduce the contribution of muscle activity that is not time-locked to stimulation in perturbational measurements (91, 92). In psychedelic states, observational complexity increases while perturbational complexity remains stable (93, 94). One possible explanation is that, under psychedelics, suppression of the alpha rhythm –a major source of signal regularity and predictability– has a stronger influence on spontaneous signal diversity than on averaged evoked responses. Another, not mutually exclusive, interpretation is that perturbational measures index the capacity for consciousness by probing the available causal interactions, whereas observational measures are more sensitive to the richness of ongoing conscious contents (93, 95). These possibilities emphasize the value of combining the two approaches rather than expecting them to vary identically across conditions.

The same framework may guide interpretation in systems whose organization is developing, unconventional, or incompletely known. In sparsely connected cortical cultures, pharmacologically increasing excitability raises observational complexity while producing only modest and inconsistent effects on perturbational complexity (96). This suggests that heightened excitability can enrich spontaneous dynamics without necessarily expanding the system’s effective causal repertoire, especially when network topology remains poorly balanced. In brain organoids, longitudinal measurements using both approaches could help distinguish changes in excitability and dynamical state from the emergence of recurrent, distributed interactions resembling those occurring in *in vivo* cortical circuits This logic may also extend beyond neural systems, including synthetic constructs such as Xenobots, which lack a canonical nervous system but have been nevertheless reported to exhibit brain-like features of spontaneous dynamics (97). Our findings caution against treating high complexity values as sufficient evidence of a particular underlying architecture and identify perturbational testing as a complementary means of constraining such inferences.

These broader implications should, however, be interpreted in light of several limitations. The model used here is deliberately simplified: networks were binary, undirected, degree-preserving, and not embedded in an anatomical space; local node dynamics were homogeneous; and the common-driver manipulation captured only one class of confound. These choices allowed us to isolate relationships between topology, dynamics, and measurement, but limit their direct generalization to biological or engineered systems. Future work should test whether the same relationships hold in weighted, directed, spatially embedded networks with heterogeneous local dynamics and biologically realistic delays.

Together, our results help close the gap between theoretical definitions of complexity and the metrics used to estimate it. Observational and perturbational approaches reveal complementary aspects of network activity, with different sensitivities to architecture, operating regime, and external influences. Establishing these relationships makes empirical findings more interpretable and provides a firmer basis for inferring functional organization both within and beyond the human brain.

## 4. Methods

### 4.1 Network definition and analysis

We used binary and undirected graphs with N=225 nodes with a fixed degree of k=24 connections per node, corresponding to a density of approximately 11%. This sparse connectivity was chosen to mimic large-scale brain networks and is comparable to empirical human structural connectomes of similar size (N=234), which exhibit mean densities on the order of 13% (98). We additionally replicated key results in the Supplementary Material using smaller networks with N=112, considering both a matched density (k=12) and different densities (k=8 and k=16), within the sparse regime. Transmission delays were assigned independently to each edge by sampling from an exponential distribution with a mean delay of 5 ms (scale parameter=5 ms), which has previously been shown to yield realistic large-scale brain dynamics in network models (99). Because our networks were defined topologically rather than in physical space, this choice represents a phenomenological approximation of heterogeneous brain transmission delays, capturing a right-skewed distribution with many short and fewer long delays (100), while isolating topological effects and preserving biologically plausible delay statistics. In the Supplementary Material, we further assessed the robustness of the main findings by repeating key analyses using alternative delay distributions, including a log-normal distribution and an empirical distribution derived from human brain data (100).

To investigate complexity as a balance between integration and differentiation, we constructed networks along a structural integration–segregation continuum, using segregation as the structural counterpart of differentiation. The first family progressed from ring lattices to increasingly random graphs, from segregation to integration; the second started from the same random graphs and progressively introduced modular organization, from integration to segregation. Ten independent realizations were generated for each family, and results are reported as averages across realizations.

#### Ring-to-random randomization

To explore the transition from segregated to integrated architectures, we implemented a rewiring procedure starting from ring lattices and progressively randomizing connections. For the main results, ten ring lattices were generated using the Watts–Strogatz model with N=225 nodes, and k=24 nearest neighbors, implemented in the NetworkX library (101). Each of the ten ring lattices was then iteratively rewired 20 times according to a degree-preserving randomization procedure based on the link-switching method (71).

From the ten original ring lattices (rewiring step=0), the subsequent 20 rewiring steps were implemented by changing the rewiring parameter *r*, distributed in a logarithmic space between 10^-3^ and 3. At each step, the network obtained in the previous step was used as input, so that the applied increment *Δr* was cumulative across steps. The *r* parameter controls the number of iterations of the link switching procedure, and a given *r* results in *r × L* rewired links, where *L* is the total number of links in the network. The result was a set of ten networks per step, resulting in 210 networks (10 initial rings × 21 steps each) ranging from ordered ring lattices to disordered random graphs.

#### Random-to-modular modularization

From the ten randomized graphs obtained in the randomization procedure, we then constructed progressively more modular networks using a novel degree-preserving clusterization algorithm, inspired by the topological overlap one (102), developed for this study. At each iteration, the algorithm selected a node at random and rewired a pair of edges to preferentially connect it with a non-neighbor sharing many common neighbors, while removing an edge associated with a less clustered local configuration. The resulting double-edge swap preserved the degree of every node and generally increased local clustering. The full algorithm is described in Supplementary Methods 1.1.

Starting from each random graph (rewiring step = 0), we generated the subsequent 20 network states by applying increasing numbers of cumulative clusterization iterations, with iteration counts logarithmically spaced between 500 and 15,000. The network obtained at each step served as the input for the next step. This procedure yielded ten networks per step, for a total of 210 networks (10 initial random graphs × 21 steps), spanning a continuum from random graphs to architectures with progressively higher clustering and stronger modular organization.

#### Network metrics

To characterize the structural properties of the generated networks, we quantified integration, segregation, and their balance using standard graph-theoretic measures. Segregation was assessed with the average clustering coefficient (13), defined as the mean probability that the neighbors of a node are also connected to each other, thereby quantifying the density of local triangular motifs. Integration was quantified using global efficiency (79), which captures how efficiently information can be exchanged across the entire network. For a given pair of nodes, efficiency is defined as the reciprocal of their shortest path length. Global efficiency is then obtained by averaging this quantity across all node pairs. These measures were computed using the *average_clustering* and *global_efficiency* functions, respectively, from the NetworkX library (101).

To evaluate the balance between network segregation and integration, we computed the small-world propensity (Muldoon et al., 2016), which jointly normalizes the observed clustering coefficient and characteristic path length (i.e., the average shortest path length between all pairs of nodes) relative to corresponding lattice and random null models. This index is bounded between 0 (no small-worldness) and 1 (maximal small-worldness) and quantifies the extent to which a network exhibits small-world structure.

### 4.2 Local node dynamics

#### Wilson-Cowan rate model

The network dynamics were modeled using a system of coupled Wilson-Cowan (WC) equations for excitatory (*E*) and inhibitory (*I*) populations at each node (103). For a given node, the population activities *r_E_* and *r_I_* evolve according to

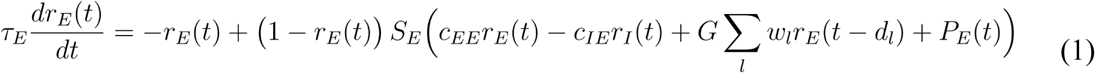

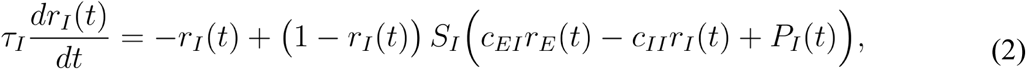

where *τ_E_* and *τ_I_* are the excitatory and inhibitory time constants, *c_XY_* the synaptic coupling strengths from population *X* to *Y*, and *G* is the global coupling scaling factor for inter-node connectivity. The connectivity weights from other nodes are denoted by *w_l_*, with corresponding transmission delays *d_l_*.

The function *S_E/I_(·)* is a sigmoidal activation function mapping synaptic inputs to firing rates:

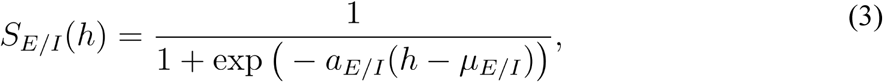

where *h* denotes the total synaptic input to the excitatory or inhibitory population, *a_E/I_*controls the slope, and *μ_E/I_* defines the input at which the function reaches its midpoint. The excitatory drive *P_E_(t)* is defined as:

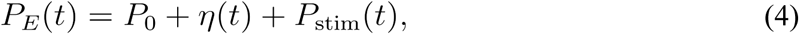

where *P_0_*is a stationary external input and *η(t)* is a stochastic drive. *P_stim_(t)* is a transient stimulation input applied exclusively to the stimulated node (square pulse, amplitude 1 a.u., duration 5 ms; *P_stim_*=0 for all other nodes). The inhibitory population receives only the stationary input, i.e.:

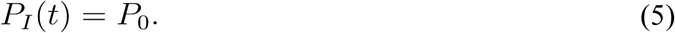

To investigate the role of independent versus shared fluctuations (e.g., common drivers) across neural populations, we extended the model by introducing a common stochastic process shared across all nodes. The stochastic drive *η(t)* entering *P_E_(t)* was defined as a normalized mixture of a node-specific independent Ornstein-Uhlenbeck (OU) process *η_I_(t)* and a common OU process *η_C_(t)* shared across all nodes. Each process evolves according to:

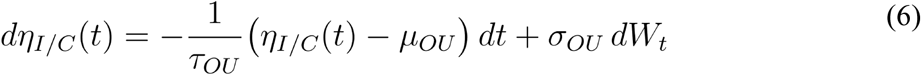

where μ is the mean, τ the correlation timescale, σ the noise amplitude, and dW_t_ a Wiener increment. Both *η_I_(t)* and *η_C_(t)* were generated with identical parameters (*μ_OU_, τ_OU_, σ_OU_*) and independent Wiener increments. The effective stochastic input is given by:

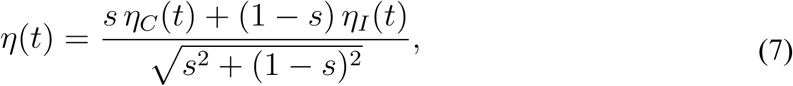

where the mixing parameter *s* ∈ [0,1] controls the relative contribution of the common driver. For *s*=0, each node receives only independent fluctuations; for *s*=1, all nodes receive identical input. The denominator acts as a normalization factor, ensuring that the variance of *η(t)* remains constant across values of *s*. Unless otherwise stated, all simulations were performed with s=0, such that each node receives only independent stochastic fluctuations.

We primarily used the default parameters of the WC model implemented in *neurolib* (104); see Supplementary Table S1). To study the bifurcation behavior of the whole network, we varied the global coupling parameter (*G*). Unless otherwise stated, we set *G=*0.1 in all simulations, placing all network architectures in a stable fixed-point regime near the supercritical Hopf stability boundary.

### 4.3 Network simulations

Simulations were performed using the *neurolib* Python library (104) with the Euler-Maruyama integration scheme and a time step of dt=0.1 ms. The output was downsampled to 1 ms after applying an anti-aliasing filter (an order-8 Chebyshev type I filter). All simulations were initialized with random initial conditions. Spontaneous activity was simulated for 63 s, with the first 3 s discarded as transient. For each network realization, one node was randomly selected for stimulation and kept fixed across 100 independent trials. The stimulation consisted of a square pulse with amplitude 1 a.u. and duration of 5 ms. At each trial, the simulation was reinitialized with random initial conditions, and after discarding the first 3 s of transient activity, the pulse was applied within a random time window between 300 and 800 ms. For analysis, the stimulus onset was centered at time 0 ms, with the pre-stimulus window defined from −300 ms to −1 ms and the post-stimulus window until 800 ms.

### 4.4 Spontaneous activity metrics

#### Mean relaxation time

To assess the proximity of each network architecture to the stability boundary, we estimated an effective relaxation time (τ) from the decay rate of damped oscillations. Since the networks are constructed with identical in-degree across nodes, at G=0.1 they converge to the same equilibrium excitatory and inhibitory firing rate (*r_e_\** and *r_i_\**, respectively). We characterized the relaxation dynamics following small deviations from this equilibrium using the raw activity traces. Starting from the equilibrium values, we independently displaced the excitatory and inhibitory firing rates of each node by +0.01, simulated the deterministic model, and extracted the excitatory activity *r_e_(t)*. For each displaced node, we identified the local maxima of the raw trace that lie above the fixed point *r_e_\**, and fitted these peak values to *r_e,peak_(t)=r_e_*+A_0_ e^−kt^*, where *r_e,peak_(t)* denotes the value of r_e_ at the identified local maximum, *A_0_* is the initial amplitude and *k* is the decay constant. The relaxation time was defined as *τ=1/k*. This procedure was repeated for 10 sampled nodes per realization, and the resulting estimates were averaged to obtain a mean *τ* (⟨*τ*⟩) for each network realization; We interpret ⟨*τ*⟩ as a numerical proxy for effective proximity to the stability boundary: larger values indicate slower return and weaker local stability, whereas smaller values indicate faster recovery and a greater dynamical margin. This empirical measure provides a tractable comparison of attractor stability across topologies, circumventing the need to compute the infinite transcendental eigenvalue spectrum of the delayed Wilson–Cowan system, a problem for which closed-form expressions for the roots do not exist.

#### Functional connectivity

Functional connectivity (FC) was estimated to capture the strength of spontaneous nodal interactions. To account for delays, we used the cross-correlation function (CCF) across time lags up to 65 ms, encompassing the maximum structural delay observed across all network topologies (i.e., 64.4 ms). FC was defined as the peak CCF value for each pair of nodes, expressed as a correlation coefficient (r) (105). From the resulting distribution of FC values, we extracted the median and the 5th and 95th percentiles, considering only the upper triangular part of the matrix.

### 4.5 Evoked responses metrics

#### Binary spatiotemporal matrix

Based on *r_E_*, significantly active nodes were identified by applying a non-parametric bootstrap-based maximal statistical procedure (Nichols and Holmes, 2001). First, evoked responses were baseline-corrected by subtracting the mean and dividing by the standard deviation of the pre-stimulus period. Next, pre-stimulus activities were bootstrapped across trials and samples. At each node and time point, the mean of the bootstrapped values (Jboot) was computed. Then, for each pre-stimulus time point, the maximum absolute Jboot across nodes was obtained. This procedure (steps 2-4) was repeated 500 times to build a null distribution. The (1−α) percentile of this distribution, with α=0.01, defined the significance threshold. Finally, a binary spatiotemporal matrix of significant activations (S) was obtained, marking samples exceeding the threshold as significant (92).

#### Active nodes

From the binary spatiotemporal activation matrix S, we computed the time course of the percentage of the summed significant activities (active nodes) across nodes and dividing by the total node count. From this time course, we extracted (i) the maximum early activation (*max_EA_*), defined as the maximum percentage of active nodes in the [0-20] ms window; (ii) the mean late activation (*mean_LA_*), defined as the average percentage of active nodes in the [450-800] ms window; and (iii) the ratio *mean_LA_ / max_EA_*.

### 4.6 Dynamical measures of complexity

To characterize the complexity of network dynamics, we analyzed two complementary classes of measures: observational metrics derived from spontaneous activity and perturbational metrics derived from stimulation-evoked responses (Supplementary Fig. S1). Observational measures quantified either signal diversity or the structure of functional interactions, including their integration–differentiation balance, whereas perturbational measures quantified the complexity, dimensionality, entropy, and temporal unpredictability of evoked spatiotemporal responses.

#### 4.6.1 Observational complexity measures

We computed five observational metrics. Lempel–Ziv complexity (LZc) and amplitude coalition entropy (ACE) quantified the diversity of ongoing amplitude patterns, whereas synchrony coalition entropy (SCE) quantified the diversity of phase-synchronization patterns (53). Neural complexity (NC) quantified multivariate statistical dependencies between complementary network partitions across spatial scales from the correlation matrix (106). Functional complexity (F_comp_) quantified the richness of the functional connectivity distribution and was intended to capture intermediate levels of functional integration and differentiation (71). Detailed formulations and normalization procedures are provided in the Supplementary Methods 1.4.

#### 4.6.2 Perturbational complexity measures

We computed four perturbational metrics from the binary spatiotemporal matrix of significant evoked activations. The perturbational complexity index (PCI) quantified the algorithmic complexity of spatiotemporal responses (92). Pattern dimensionality measured the effective number of principal components required to explain the evoked-response variance (107). Pattern entropy quantified the diversity of spatial and temporal activation patterns (108), and entropy rate quantified the temporal unpredictability of node-wise activation sequences (109). Detailed formulations, temporal coarse-graining, and bias correction are provided in the Supplementary Methods 1.5.

### 4.7 Statistical analysis

Statistical analyses were performed in Python. Repeated-measures ANOVA was used to test effects across rewiring steps, Spearman’s ρ was used for correlation analyses between measures, and linear regression was used to assess explanatory relationships. Shaded regions denote the standard error of the mean (SEM). To quantify the sensitivity of each complexity metric to common drivers, we computed for each network realization the mean absolute percentage deviation from the *s=*0 baseline across all tested common driver strengths (avg. % deviation from baseline). Formally, this is defined as the mean of (|M(s_i_) − M(0)| / M(0)) × 100 over all N common driver strength values s_i_ > 0, where M(0) is the metric value in the absence of common driver.

## Data, Materials, and Software Availability

The code used to perform the simulations and reproduce the results presented in this article will be made publicly available upon publication at https://github.com/gianlucagag/NetworkComplexity.

## Acknowledgments

The authors thank Michele Colombo and Marta Porro for their comments and suggestions on the manuscript and figures. This work was financially supported by the following entities: ERC-2022-SYG Grant number 101071900 neurological mechanisms of injury and sleep-like cellular dynamics (NEMESIS); Tiny Blue Dot Foundation; Canadian Institute for Advanced Research (CIFAR), Canada.

## Author contributions

G.G., A.C., T.N., D.M., R.C., L.D.P., S.S., and M.M. designed research; G.G. performed simulations; G.G. analyzed data; G.G. organized results; G.G., T.N., L.D.P, and R.C. designed figures; G.G. wrote the first draft; All authors revised and approved the paper.

