## Supplementary information for "Inferring brain structural complexity from dynamical measures"

#### Table of Contents


1. **Supplementary Methods**
    - 1.1 Degree-preserving clusterization algorithm
    - 1.2 Wilson-Cowan model with spike-frequency adaptation
    - 1.3 Network amplitude fluctuations
    - 1.4 Observational complexity measures
    - 1.5 Perturbational complexity measures
2. **Supplementary Figures**
    - Figure S1. Schematic overview of the dynamical measures of complexity used in this study
    - Figure S2. Collective network fluctuations near the critical operating point
    - Figure S3. Rewiring-induced changes in spontaneous functional connectivity and evoked responses
    - Figure S4. Complexity measures across network densities
    - Figure S5. Input–output characterization and perturbational complexity across stimulation amplitudes
    - Figure S6. Effect of transmission-delay distribution on observational and perturbational complexity across rewiring
    - Figure S7. Correlation among structural, functional and dynamical measures
    - Figure S8. Identification of highly clustered networks in the modularization process
    - Figure S9. Perturbational complexity measures as a function of global and local network properties
    - Figure S10. PCI partially reflects the duration of the evoked activity
    - Figure S11. Perturbational complexity reflects small-world organization beyond differences in relaxation dynamics
3. **Supplementary Tables**
    - Table S1. Parameters of the standard Wilson-Cowan model
    - Table S2. Parameters of the Wilson-Cowan model with spike-frequency adaptation
4. **Supplementary Notes**
    - 4.1 Perturbational complexity reflects small-world organization beyond local attractor stability
    - 4.2 Local perturbation gain, network amplification, and attractor stability
5. **Supplementary References**

### 1. Supplementary Methods

#### 1.1 Degree-preserving clusterization algorithm

We developed a stochastic degree-preserving rewiring algorithm to increase local clustering and promote modular organization while preserving the degree of every node. The algorithm was designed for simple, undirected, unweighted graphs represented by a symmetric binary adjacency matrix. At each iteration, a node $i$ was selected uniformly at random.
Let $\mathcal{N}(i)$ denote the neighbors of $i$, and let $\overline{\mathcal{N}(i)}$ denote the nodes that are neither $i$ nor neighbors of $i$. For any two nodes $u$ and $v$, we define their common-neighbor count as

$$\mathrm{CN}(u,v) = |\mathcal{N}(u) \cap \mathcal{N}(v)|.$$

Among the non-neighbors of $i$, the algorithm selected a node $j$ with the largest value of $\mathrm{CN}(i,j)$. Among the neighbors of $i$, the algorithm selected a node $k$ with the smallest value of $\mathrm{CN}(i,k)$. Thus, the proposed rewiring removes the edge $(i,k)$ and adds the edge $(i,j)$, preferentially connecting $i$ to a node with which it already shares many neighbors.

To complete a degree-preserving double-edge swap, the neighbors of $j$ were ordered according to increasing $\mathrm{CN}(j,l)$. The algorithm then selected the first candidate $l$ in this order that satisfied $(k,l) \notin E$, where $E$ is the current edge set. The edges $(j,l)$ and $(i,k)$ were removed, and the edges $(i,j)$ and $(k,l)$ were added. The swap was accepted only when $i$, $j$, $k$, and $l$ were four distinct nodes. If no admissible $l$ existed, or if the four-node distinctness condition was violated, the iteration was discarded and the graph was left unchanged.

The accepted edge replacement can be written as

$$\{(i,k),(j,l)\} \longrightarrow \{(i,j),(k,l)\}.$$

Because each of the four nodes loses one incident edge and gains one incident edge, the degree sequence is preserved exactly. The procedure is heuristic rather than an optimization of a globally defined objective: it preferentially replaces an edge incident to a node sharing few common neighbors with an edge connecting nodes that share many common neighbors. This local rule is intended to increase the number of triangular motifs and, consequently, the average clustering coefficient. The algorithm does not guarantee that clustering increases at every accepted iteration, nor does it impose a target modularity value. Its effect on network organization was therefore evaluated using the graph-theoretic measures described in the main Methods section.

**Algorithm 1. Degree-preserving clusterization rewiring**

**Input:** Binary adjacency matrix $A \in \{0,1\}^{n \times n}$, number of iterations $I$
**Output:** Rewired adjacency matrix $A'$

1. $G \leftarrow \text{GraphFromMatrix}(A)$
2. For $t = 1, \dots, I$:
3. $\quad v \leftarrow \text{UniformRandomInteger}(0, n-1)$
4. $\quad C \leftarrow \text{Neighbors}(G, v)$
5. $\quad U \leftarrow V(G) \setminus \big(C \cup \{v\}\big)$
6. If $|C| = 0$ or $|U| = 0$, continue
7. For each $c \in C$, compute

$$q_c \leftarrow |\text{Neighbors}(G, v) \cap \text{Neighbors}(G, c)|$$

8. For each $u \in U$, compute

$$q_u \leftarrow |\text{Neighbors}(G, v) \cap \text{Neighbors}(G, u)|$$

9. $\quad j \leftarrow \arg\max_{u \in U} q_u$

▷ candidate node to add

10. $\quad k \leftarrow \arg\min_{c \in C} q_c$

▷ candidate node to remove

11. $\quad C_j \leftarrow \text{Neighbors}(G, j)$
12. If $|C_j| = 0$, continue
13. Order the nodes $l \in C_j$ by increasing

$$q_l' \leftarrow |\text{Neighbors}(G, j) \cap \text{Neighbors}(G, l)|$$

14. $\quad l \leftarrow$ first ordered node satisfying $(k, l) \notin E(G)$
15. If no such node exists, continue
16. If $|\{v, j, k, l\}| < 4$, continue
17. Remove edges $(v, k)$ and $(j, l)$
18. Add edges $(v, j)$ and $(k, l)$
19. End for
20. $A' \leftarrow \text{MatrixFromGraph}(G)$
21. Return $A'$

##### 1.2 Wilson-Cowan model with spike-frequency adaptation

We implemented in The Virtual Brain (TVB) a Wilson-Cowan rate model with spike-frequency adaptation current (WCA) as in (1, 2). This model reproduces key features of Up and Down state transitions arising from the interplay between adaptation, recurrent excitation, external inputs, and noise. The WCA comprises three state variables representing the excitatory firing rate $r_e$, inhibitory firing rate $r_i$, and adaptation current $a$. Two independent Ornstein-Uhlenbeck (OU) noise processes, $u_e$ and $u_i$, act on the excitatory and inhibitory populations, respectively. The population firing rates evolve according to:

$$\tau_e \frac{dr_e}{dt} = -r_e + R_e(w_{ee}r_e - w_{ie}r_i + c\sum_k W_{jk}r_e^k[t-\tau_{jk}] + I_e - a + u_e), \quad (1)$$

$$\tau_i \frac{dr_i}{dt} = -r_i + R_i(w_{ei}r_e - w_{ii}r_i + I_i + u_i), \quad (2)$$

where $R_e$ and $R_i$ are power-law input-output functions (3) defined as:

$$R_{e/i,\infty}(x) = k_{e/i}\,[x - h_{e/i}]_+^n, \quad (3)$$

with scaling factors $k_e$ and $k_i$, thresholds $h_e$ and $h_i$, and exponent $n$. The internal coupling terms include recurrent weights $w_{ee}$ (excitatory-to-excitatory), $w_{ei}$ (excitatory-to-inhibitory), $w_{ie}$ (inhibitory-to-excitatory), and $w_{ii}$ (inhibitory-to-inhibitory neurons); the network coupling, which targets the excitatory population, is given by $c\sum_k W_{jk}r_e^k[t\text{-}\tau_{jk}]$, where $W_{jk}$ is the connection weight from presynaptic node $k$ to postsynaptic node $j$, and $\tau_{jk}$ represents the time delay between $j$ and $k$. The term $c$ is the global coupling factor that scales all the connection weights. Thus, the effective excitatory coupling strength between $j$ and $k$ is given by the product of $W_{jk}$ and $c$, and we denote it by $C_{jk}$. $r_e^k$ is the firing rate of node $k$, and $I_e$ and $I_i$ are external inputs to excitatory and inhibitory populations, respectively. The slow adaptation current $a$ implements spike-frequency adaptation through:

$$\tau_a \frac{da}{dt} = -a + A_\infty(r_e), \quad (4)$$

where

$$A_\infty(r_e) = \frac{g}{1 + e^{-a_a(r_e - \mu_a)}}, \quad (5)$$

is a sigmoid function of the excitatory firing rate with gain $a_a$, threshold $\mu_a$, and strength $g$ (denoted as b in the original article). $\tau_a$ is the time constant of the adaptation dynamics. The noise processes follow the OU dynamics:

$$\tau_{ou} \frac{du_{e/i}}{dt} = -u_{e/i} + \sigma_{ou}\xi_{e/i}(t), \quad (6)$$

where $\xi_{e\backslash i}(t)$ are independent Wiener processes with mean $\mu_{e/i}$, amplitude $\sigma_{ou}$, and time constant $\tau_{ou}$. The parameters used for the WCA simulations are listed in Table S2. Equations were integrated using the Heun stochastic method in TVB with a time step of 0.5 ms.

#### 1.3 Network amplitude fluctuations.

To characterize slow collective fluctuations in network activity in a subset of analyses, simulations were extended to 250 s to ensure reliable estimation of infraslow dynamics. Although metastability is commonly quantified as the temporal standard deviation of the Kuramoto order parameter (e.g., (4)), this measure relies on instantaneous phase synchronization and is more difficult to interpret in systems with fast oscillations and non-negligible transmission delays. Instead, collective dynamics were characterized via the global amplitude signal. For each node, $r_E$ was transformed into amplitude using the Hilbert transform, and node amplitudes were averaged to obtain a global amplitude time series. Its temporal standard deviation was used as a measure of collective fluctuation strength. To test whether these fluctuations depended on coordinated interactions, a surrogate distribution was generated (100 realizations). For each surrogate, node activities were independently circularly shifted by a random amount (≥1 s), preserving temporal structure and amplitude statistics while disrupting temporal alignment. The global amplitude was recomputed for each surrogate, yielding a null distribution of fluctuation strengths. The observed value was expressed as a z-score relative to this distribution.

#### 1.4 Observational complexity measures

**Lempel-Ziv complexity.** Lempel-Ziv complexity (LZc) was used to quantify the diversity of spatiotemporal patterns and implemented following (5). After applying the Hilbert transform, the amplitude signal was binarized using the mean amplitude as a threshold. This yielded a binary spatiotemporal activity matrix, which was spatially concatenated before applying the LZ76 algorithm (6). Because spatial concatenation makes LZc potentially sensitive to node ordering, node labels were randomly permuted before concatenation. This removed any privileged ordering present in the ring networks, where adjacent node indices also correspond to topological neighbors, and therefore prevented random networks from being penalized simply because their node ordering carries no comparable structural information. Normalization was performed by dividing the raw value by the LZc of the same data temporally shuffled.

**Coalition entropy.** Similarly to LZc, Amplitude coalition entropy (ACE) and Synchrony coalition entropy (SCE) were computed as in (5) to capture the richness of spatiotemporal activation patterns and phase synchronization patterns, respectively. ACE is the Shannon entropy, over time, of the set (coalition) of active nodes, based on the same binarization procedure used for LZc; ACE values were normalized against temporally shuffled data. SCE is the Shannon entropy, over time, of synchrony coalitions obtained by binarizing, for each reference node, all other nodes according to whether their instantaneous Hilbert phase difference with the reference node was less than 0.8 radians. The coalition entropy for each reference node was normalized by the entropy of randomly generated binary coalitions in which each entry was independently set to 1 with probability 0.5, representing the theoretical upper bound. SCE values were then averaged across reference nodes.

**Neural complexity.** Neural complexity (NC) was computed from the correlation matrix following (6). For each bipartition of the network into two complementary subsets, mutual information was estimated from the determinants of the full correlation matrix and of the two corresponding submatrices, as: $MI = 0.5 \times [\log(\det(C_A)) + \log(\det(C_B)) - \log(\det(C))]$, where $C$ is the full correlation matrix and $C_A$ and $C_B$ are the correlation submatrices corresponding to the two subsets. Bipartitions were evaluated across all subset sizes from 1 to $\lfloor N/2 \rfloor$. For each partition size, 100 bipartitions were randomly sampled and their mutual information values were averaged. NC was then defined as the sum of these average values across partition sizes. Thus, NC quantifies multivariate statistical dependencies between complementary subsets of the network across spatial scales.

**Functional complexity.** Functional complexity ($F_{comp}$) was computed as described in (7). $F_{comp}$ quantifies the balance between functional integration and differentiation by measuring the richness of interactions in the FC matrix (derived as described above). Low functional complexity corresponds to narrow FC distributions, reflecting either statistical independence ($\langle FC \rangle \approx 0$; differentiation) or global synchrony ($\langle FC \rangle \approx 1$; integration). Higher values arise when intermediate states yield broad FC distributions. $F_{comp}$ was quantified as the sum of absolute differences between the FC distribution and a uniform distribution across m bins.

##### 1.5 Perturbational complexity measures

**Perturbational complexity index.** The Perturbational Complexity Index (PCI) was computed as the Lempel-Ziv complexity of the binary spatiotemporal matrix S, normalized by its source entropy, as described in Casali et al. (2013). The nodes in S were sorted in descending order of their total activity during the post-stimulus period prior to compression. The LZ76 algorithm was then applied to the spatially concatenated binary sequence obtained from the sorted matrix. The resulting complexity value was normalized by the source entropy of S, defined as $H = -p \cdot \log_2(p) - (1-p) \cdot \log_2(1-p)$, where $p$ is the overall fraction of active entries in the matrix.

**Pattern dimensionality.** To quantify the effective number of spatial covariance modes in the evoked response, we applied principal component analysis to the spatiotemporal matrix S. Pattern dimensionality was defined as D=k/N, where k is the smallest number of principal components required to explain 95% of the total variance, and N is the number of nodes (8). D ranges from 1/N, when a single mode dominates, to 1, when variance is spread across modes.

**Pattern entropy.** The richness of binary network activation patterns in the evoked response was quantified by computing the Shannon entropy of the pattern distribution from the binary spatiotemporal matrix S, following (9). Whereas Shew et al. (2011) computed spatial pattern entropy across repeated stimulus trials, here we evaluated spatial patterns across time points and temporal patterns across nodes, to characterize the spatial and temporal richness of statistically significant evoked responses. Spatial pattern entropy $H_{space}$ was thus defined as the Shannon entropy over the distribution of unique spatial activity patterns (i.e., binary vectors indicating which of the N nodes were active at each time point) observed across time points; temporal pattern entropy $H_{time}$ was defined as the Shannon entropy over the distribution of unique temporal activity patterns (binary vectors indicating at which time points each node was active), observed across nodes. The overall evoked pattern entropy was defined as their average: $\langle H \rangle = (H_{space} + H_{time}) / 2$.

Prior to entropy computation, S was coarse-grained in time using a majority-vote rule over non-overlapping windows of width 10 ms, assigning state 1 to a bin if a node was active in at least half of the 10 time points within that window. To correct for finite-sample bias, entropy was estimated by randomly subsampling the pattern set at fractions f from 0.1 to 1.0 in steps of 0.1, repeating 10 times per fraction, and fitting the average H(f) with the quadratic function (Magri et al., 2009; Shew et al., 2011): $H(f) = H_0 - a/(f \cdot T) - b/(f \cdot T)^2$, where T is the total number of patterns and the fit parameter $H_0$ is the bias-corrected estimate reported in all analyses.

**Entropy rate.** The temporal unpredictability of the binary activation sequences in the evoked response was characterized using the entropy rate, estimated per node from the rows of the binary spatiotemporal matrix S. For each node and each history length k from 1 to 5, the joint Shannon entropy H(k) was computed as the Shannon entropy over the distribution of all k-length subsequences (k-histories) observed in that node's binary activation time series. The entropy rate was then estimated as the slope of the linear fit of H(k) as a function of k, using the *entropy_rate* function of the *NeuroKit2* library (10). This slope quantifies the average amount of new information generated per time step, given knowledge of the past k states. The entropy rate was averaged across nodes; nodes with no significant activations were excluded from the mean unless otherwise stated, in which case they contributed a rate of zero.

#### 2. Supplementary Figures

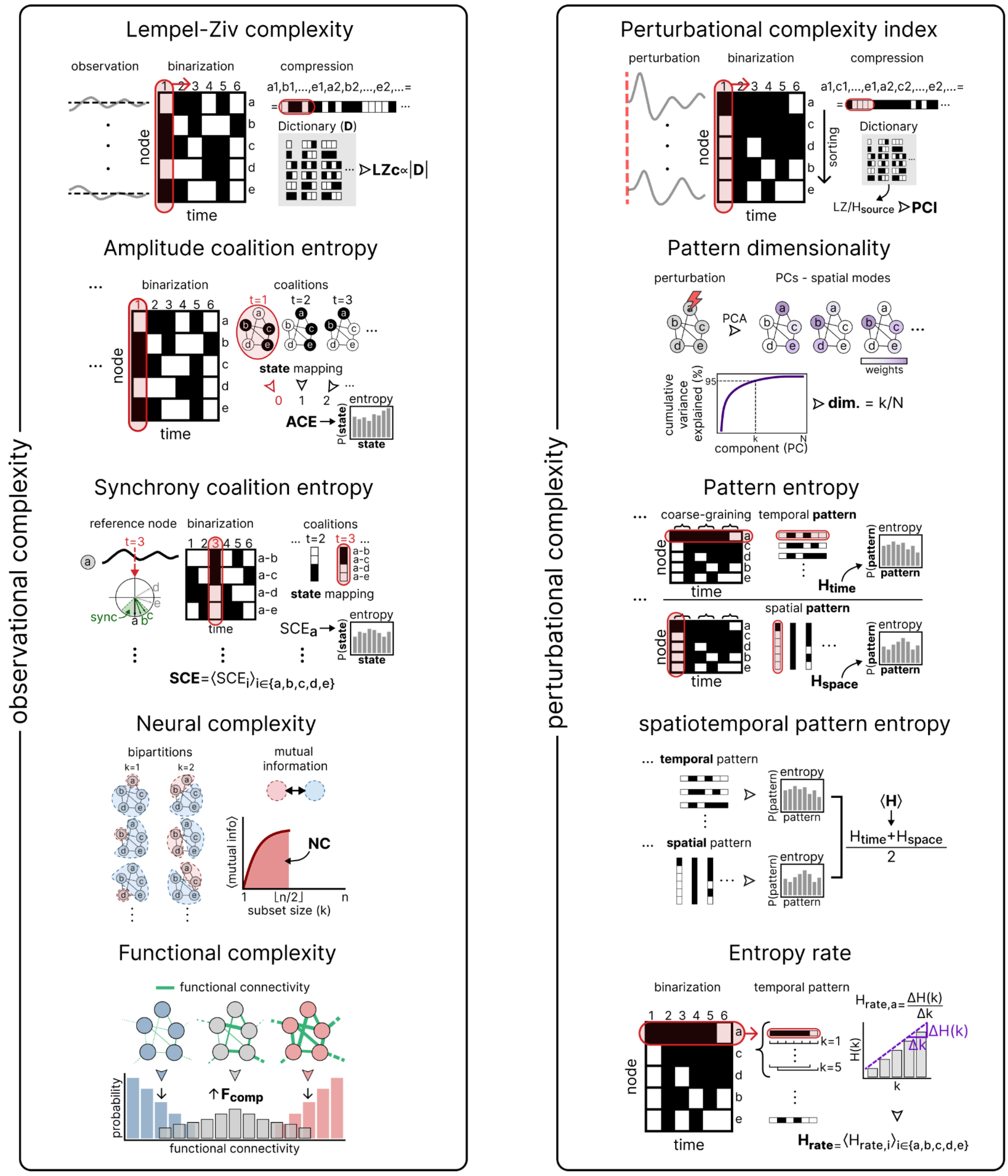


**Figure S1. Schematic overview of the dynamical measures of complexity used in this study**. Metrics are grouped into observational metrics derived from spontaneous activity (left) and perturbational metrics derived from stimulation-evoked responses (right). The diagrams summarize the main processing steps underlying each measure and are intended as conceptual illustrations rather than formal definitions.

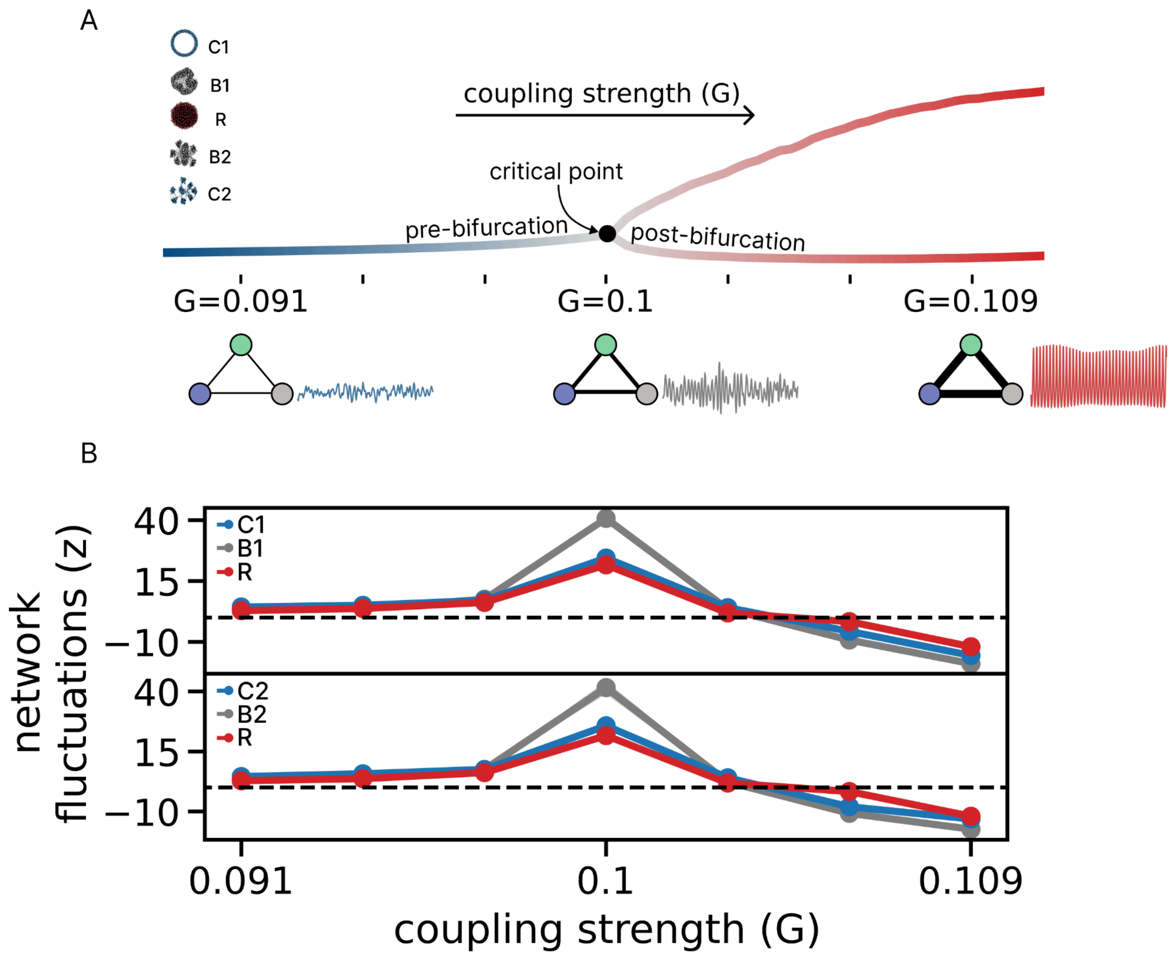


**Figure S2. Collective network fluctuations near the critical operating point. (A)** Regime sweep of C1, B1, R, B2, and C2 networks obtained by varying global coupling (G) around the critical point (G=0.091 – 0.109, step = 0.003). The bifurcation diagram for C1 is shown for illustration, with representative activity traces at pre-bifurcation (G=0.091), near-bifurcation (G=0.1), and post-bifurcation (G=0.109) regimes. **(B)** Strength of coordinated global amplitude fluctuations as a function of global coupling (G), quantified from the temporal standard deviation of the global amplitude signal (Supplementary Methods 1.3). Values are expressed as z-scores relative to circular-shift surrogate distributions that preserve individual node dynamics while disrupting temporal coordination. Fluctuation strength peaked near the critical operating point, consistent with enhanced metastable dynamics close to the instability.

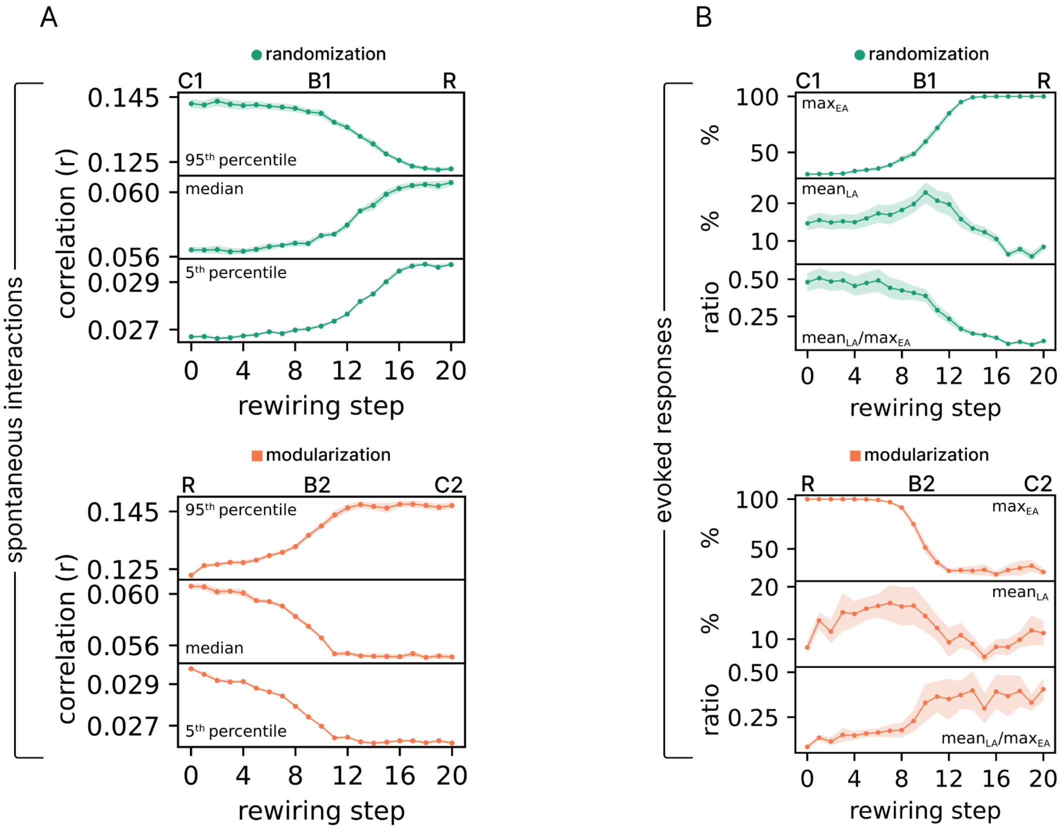


**Figure S3. Rewiring-induced changes in spontaneous functional connectivity and evoked responses. (A)** Evolution of FC statistics across rewiring steps. For each step, the distribution of correlations was summarized by the 5th, 50th (median), and 95th percentiles. Randomization (top; green curves) shifted the FC distribution toward higher median and 5th percentile values, while reducing the 95th percentile, whereas modularization (bottom; orange curves) increased the 95th percentile consistently with the emergence of a pronounced right tail. **(B)** Evolution of evoked response metrics across rewiring steps. For each family, the maximum early activation ($max_{EA}$), the mean late activation ($mean_{LA}$), and their ratio ($mean_{LA}/max_{EA}$) were quantified. Randomization (top; green curves) progressively strengthened the early network response, as shown by the increase in $max_{EA}$, but reduced the late response, as shown by the decrease in $mean_{LA}$. In contrast, modularization (bottom; orange curves) had the opposite effect: early recruitment weakened, with a reduction in $max_{EA}$, whereas activity was relatively better sustained in the activated nodes, leading to an increase in the $mean_{LA}/max_{EA}$ ratio. Shaded areas denote SEM across network realizations (10 per step).

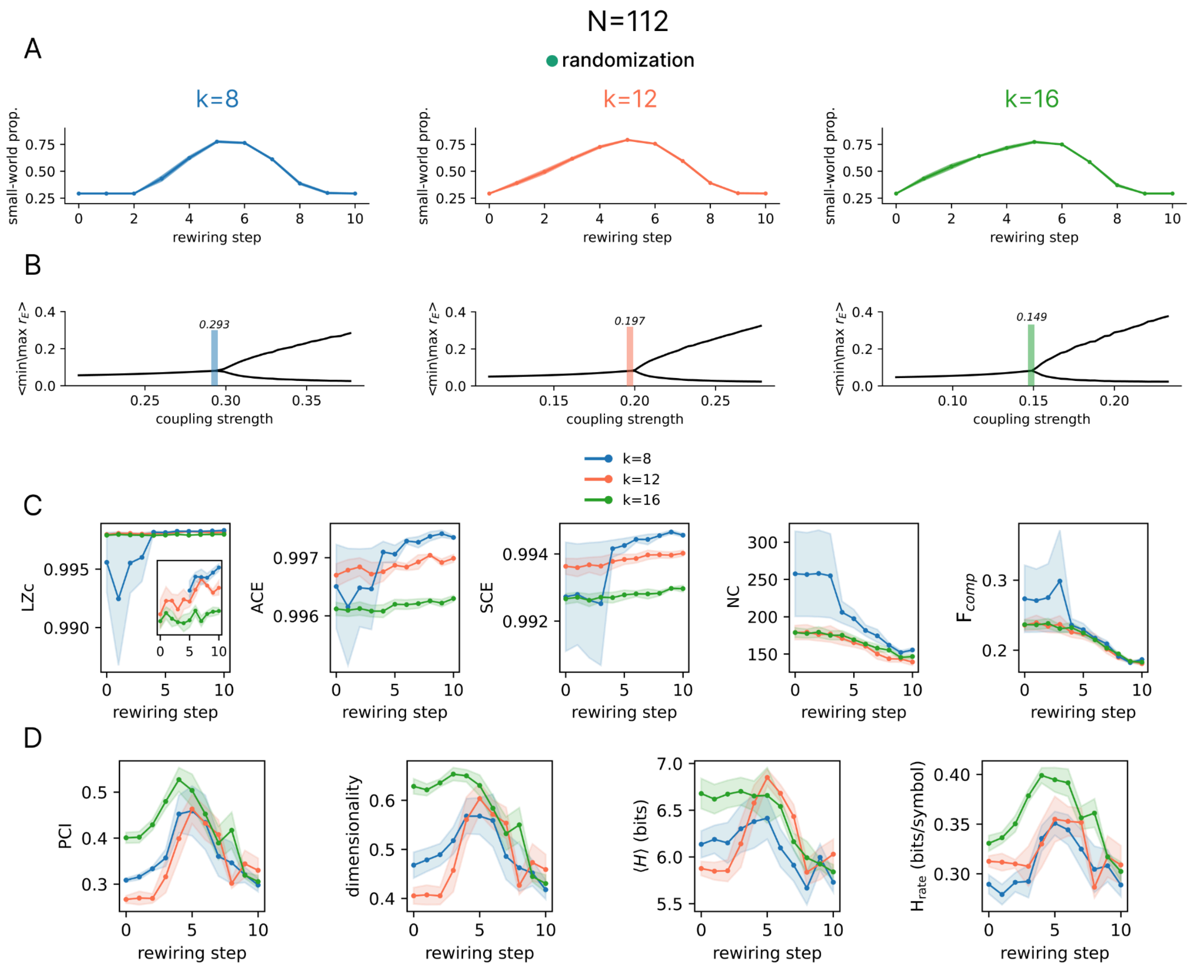


**Figure S4. Complexity measures across network densities. (A)** Small-world propensity across the randomization process for networks with N=112 nodes and k=8, 12, or 16 connections per node. Across densities, small-world propensity peaked at intermediate rewiring steps. **(B)** Dynamical regime as a function of coupling strength for each network density. Colored vertical lines indicate the coupling strengths used for simulations k=8: 0.293; k=12: 0.197; k=16: 0.149, selected to place the networks near the oscillatory instability. **(C)** Observational complexity measures (LZc, ACE, SCE, NC, and $F_{comp}$) across rewiring steps. **(D)** Perturbational complexity measures (PCI, dimensionality, ⟨H⟩, and $H_{rate}$) across rewiring steps. Lines and shaded areas indicate mean ± SEM across five network realizations.

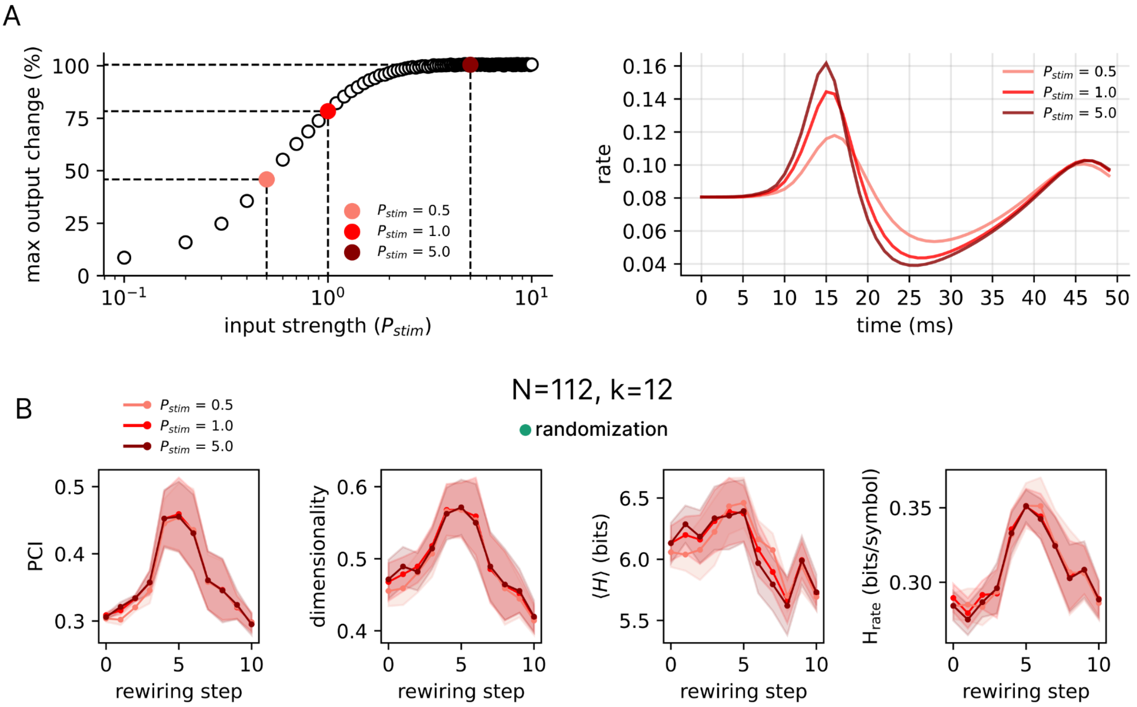


**Figure S5. Input-output characterization and perturbational complexity across stimulation amplitudes.** **(A)** Input-output characterization of the perturbation. Left, percentage change in network output as a function of stimulation strength $P_{stim}$. To quantify the network response beyond the directly stimulated node, output change was defined as the percentage change in the maximum firing rate reached across all non-stimulated nodes and time points after stimulation, relative to the corresponding maximum during the pre-stimulus period. Highlighted points indicate the three stimulation amplitudes used in the subsequent analysis $P_{stim}$=0.5, 1.0, 5.0, spanning weak to near-saturating network responses. $P_{stim}$=1.0 was used in the main analyses. Right, corresponding evoked response profiles for the three stimulation amplitudes. **(B)** Perturbational complexity measures (PCI, dimensionality, ⟨H⟩, and $H_{rate}$) across the randomization process for each stimulation amplitude, computed for networks with N=112 nodes and k=12 connections per node. Lines and shaded areas indicate mean ± SEM across five network realizations.

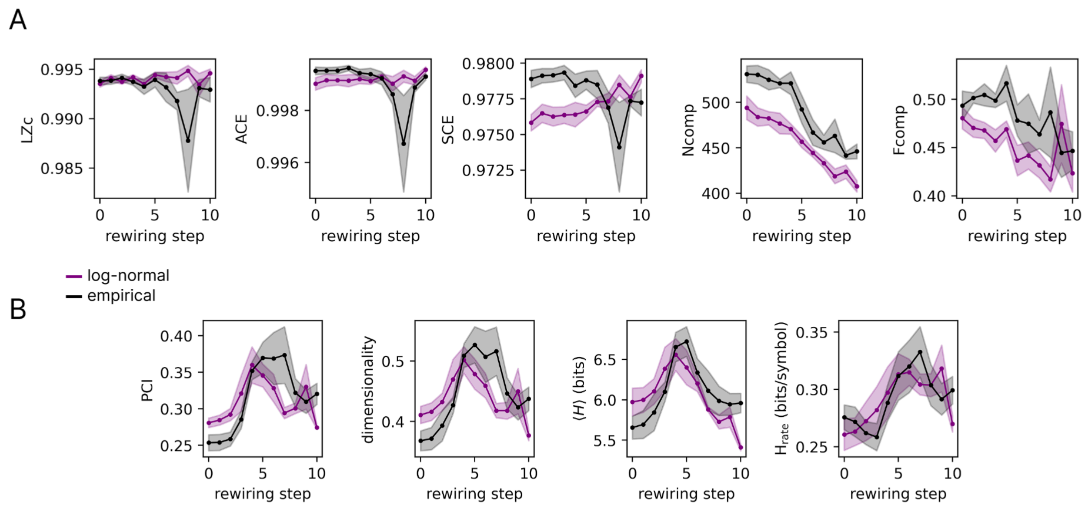


**Figure S6. Effect of transmission-delay distribution on observational and perturbational complexity across rewiring.** Complexity measures were computed across the randomization process for networks with N=112 nodes and k=12 connections per node, using either a log-normal distribution of transmission delays (purple) or an empirical delay distribution derived from human brain data (black) from (11). **(A)** Observational complexity measures: Lempel–Ziv complexity (LZc), amplitude coalition entropy (ACE), synchrony coalition entropy (SCE), neural complexity (NC), and functional complexity ($F_{comp}$). **(B)** Perturbational complexity measures: perturbational complexity index (PCI), pattern dimensionality, mean spatiotemporal entropy ($\langle H \rangle$), and entropy rate ($H_{rate}$). Lines indicate mean values across 5 network realizations and shaded areas indicate SEM.

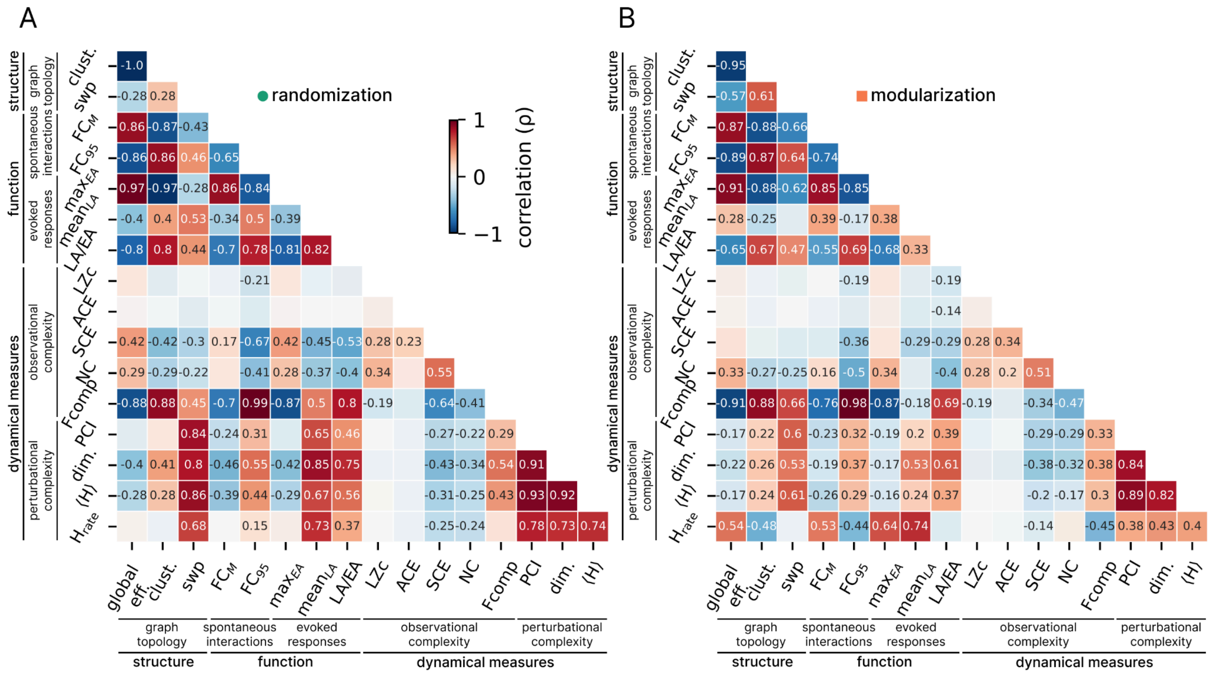


**Figure S7. Correlation among structural, functional, and dynamical measures.** Pairwise Spearman correlations (ρ) across rewiring steps are shown for **(A)** randomization and **(B)** modularization processes. Structural measures (graph-theoretical metrics) include global efficiency (eff.), average clustering coefficient (clust.), and small-world propensity (swp). Functional measures of spontaneous interactions include the median of the FC distribution ($FC_M$) and its 95th percentile ($FC_{95}$). Functional measures of evoked responses include the maximum early activation ($max_{EA}$), the mean late activation ($mean_{LA}$), and their ratio (LA/EA). Dynamical measures comprise observational complexity metrics (LZc, ACE, SCE, NC, $F_{comp}$) and perturbational complexity measures (PCI, dim., ⟨H⟩, and $H_{rate}$). Color indicates correlation strength and direction, with red denoting positive and blue denoting negative correlations. Only correlations with p-values $< 0.05$ are displayed.

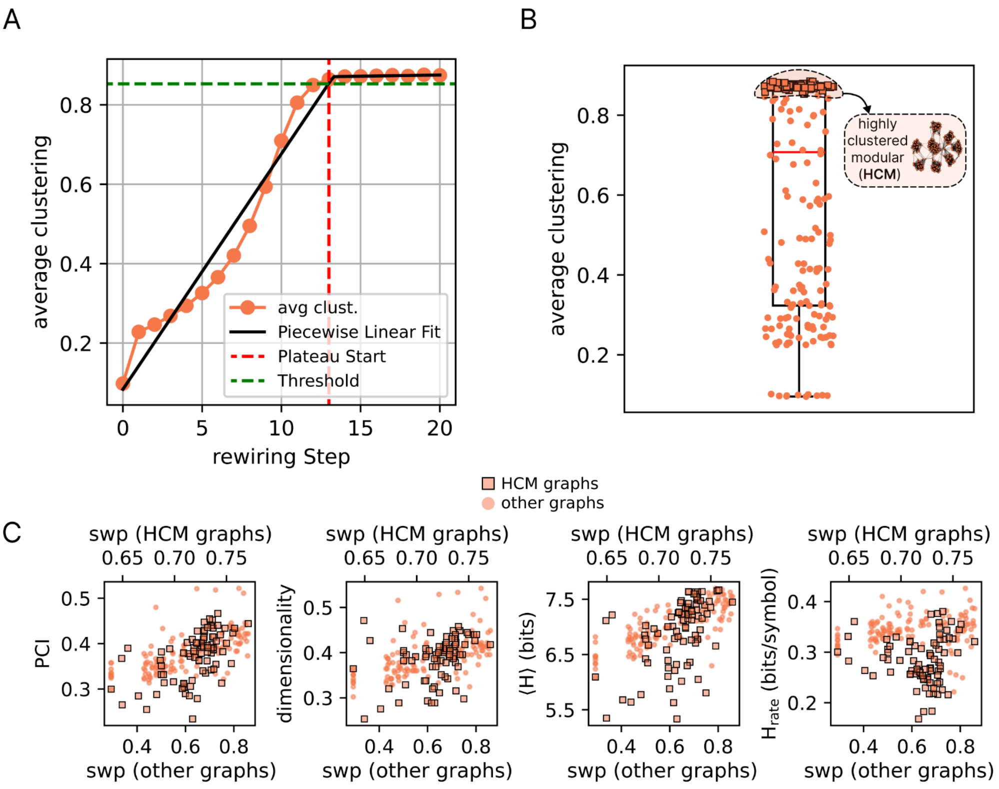


**Figure S8. Identification of highly clustered networks in the modularization process. (A)** The threshold for identifying the subgroup of highly clustered networks was derived from the evolution of average clustering during the modularization process. The mean clustering trajectory across rewiring steps (orange circles) was fitted with a two-segment piecewise linear model (black line), capturing the initial rise and the subsequent plateau. The breakpoint of this fit (red dashed line) marked the onset of the plateau, and all networks sampled at steps beyond this point were considered part of the plateau regime. To define a permissive threshold within this regime, we computed the mean clustering value across plateau networks and subtracted three standard deviations (green dashed line). **(B)** Distribution of clustering values across all networks (10 realizations for each of the 21 rewiring steps). Networks exceeding the threshold are highlighted with orange square markers and were classified as highly clustered. **(C)** Scatter plots showing perturbational complexity measures (PCI, dim., ⟨H⟩, and Hrate) as a function of small-world propensity across rewiring steps of the modularization process. Highly clustered modular (HCM) networks are highlighted as square markers; the rest of the graphs are highlighted as circle markers. Spearman correlation coefficients are reported separately in the main Fig. 3D.

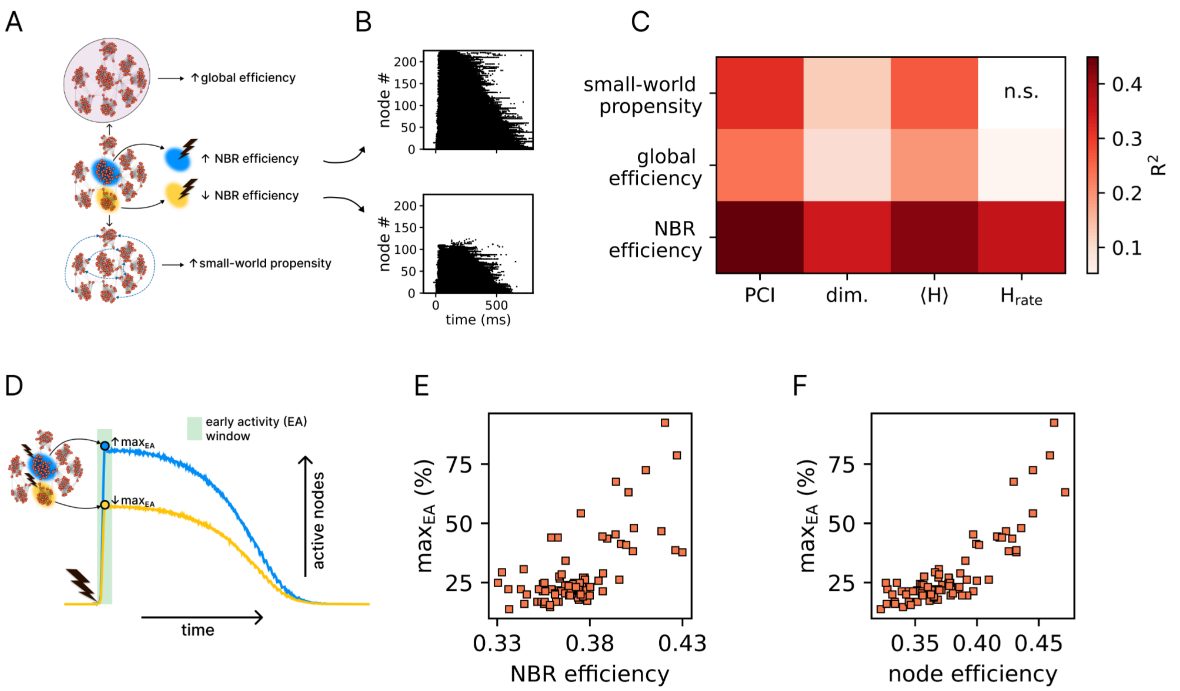


**Figure S9. Perturbational complexity measures as a function of global and local network properties. (A)** Schematic illustrating how structural properties differ across the highly clustered modular networks. Networks vary in small-world propensity and global efficiency, but also in the neighborhood (NBR) efficiency of the stimulated node. NBR efficiency was computed as the average efficiency of the stimulated node and its first-degree neighbors with respect to the whole network; accordingly, stimulation may target a node embedded in a hub-like neighborhood (high NBR efficiency, blue node) or in a peripheral neighborhood (low NBR efficiency, yellow node). **(B)** Representative matrix of significant node activations (with nodes sorted from most to least active) for two networks with contrasting NBR efficiency: stimulation of a node belonging to a hub-like cluster (top) elicits a richer spatiotemporal activation pattern compared to stimulation of a peripheral node (bottom). **(C)** Heatmap of the coefficient of determination ($R^2$) from linear regression of perturbational complexity measures onto small-world propensity, global efficiency, and NBR efficiency within the highly clustered subgroup. NBR efficiency accounts for the largest proportion of variance across all metrics; n.s. indicates a non-significant fit for $H_{rate}$. **(D)** Schematic illustration of the dependence of the maximum early activation ($max_{EA}$) on stimulation target: stimulation of a node embedded in a high-efficiency neighborhood produces a larger early network response, whereas stimulation of a node in a low-efficiency neighborhood produces a weaker one. **(E)** Scatterplot showing the positive association between NBR efficiency and $max_{EA}$ across stimulated networks ($\rho$=0.53, p=$2\times10^{-7}$) **(F)** Same analysis as in (E), but considering only the efficiency of the stimulated node ($\rho$=0.75, p=$2\times10^{-16}$).

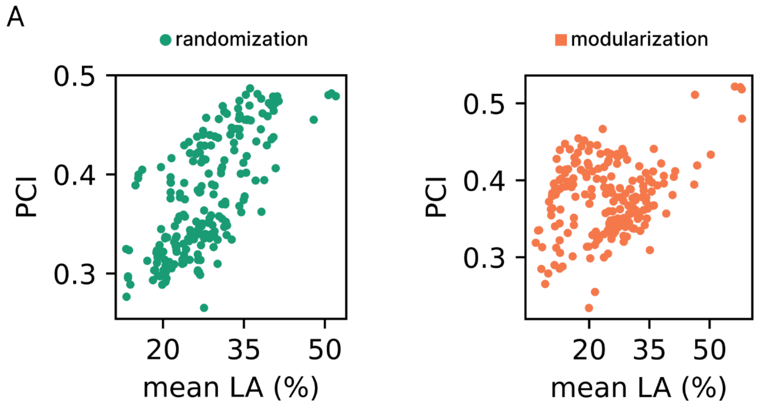


**Figure S10. PCI partially reflects the duration of the evoked activity. (A)** PCI correlates significantly with $mean_{LA}$ (randomization: ρ=0.65, p=7×10$^{-27}$; modularization: ρ=0.2, p=0.004). However, the association is moderate in the modularization process.

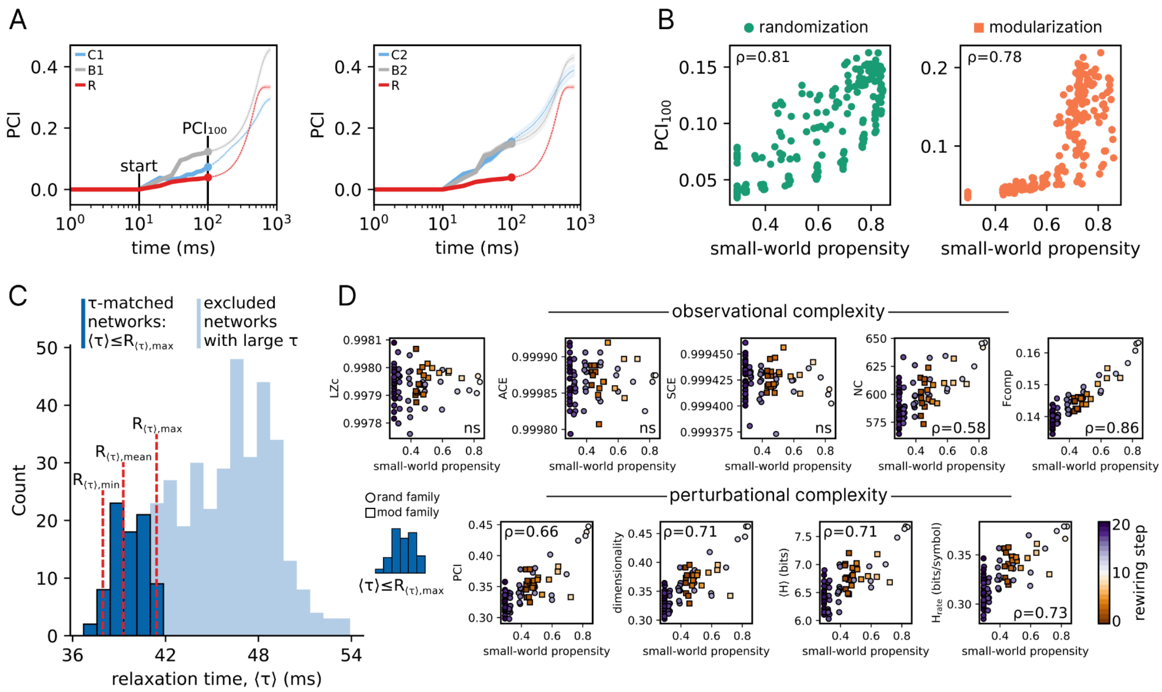

**Figure S11. Perturbational complexity reflects small-world organization beyond differences in relaxation dynamics. (A)** Time-resolved PCI following stimulation for representative clustered (C1/C2; light blue), balanced (B1/B2; gray), and random (R; red) architectures. PCI at 100 ms post-stimulation ($PCI_{100}$) was used to quantify early perturbational complexity. Shaded areas indicate mean ± SEM across network realizations. The vertical marker at 10 ms indicates the onset of PCI computation. **(B)** $PCI_{100}$ increases with small-world propensity in both the randomization (green) and modularization (orange) families. Reported $\varrho$ indicate significant Spearman rank correlations (all $p<10^{-42}$); **(C)** Selection of the relaxation time-matched subset ($\tau$-matched networks). Dark blue bars denote the τ-matched subset retained for the analysis, defined by $\langle\tau\rangle \leq R_{\langle\tau\rangle,max}$, where $R_{\langle\tau\rangle,max}$=41.4 is the largest mean relaxation time ($\langle\tau\rangle$) among the random networks (R). The retained subset comprised all 10 R networks and 71 $\tau$-matched networks from the randomization and modularization families. The mean, minimum, and maximum $\langle\tau\rangle$ of the R networks are indicated by dashed red lines. Light blue bars show networks excluded from the matched analysis because of large $\langle\tau\rangle$. **(D)** Associations between small-world propensity and observational complexity (upper row) and perturbational complexity (lower row) in the $\tau$-matched subset. Circles and squares denote networks from the randomization and modularization families, respectively; color indicates rewiring step. Reported $\varrho$ values indicate significant Spearman rank correlations (all $p<10^{-7}$); "ns" denotes non-significant correlations.

#### 3. Supplementary Tables

**Table S1. Parameters of the standard Wilson-Cowan model.** The table lists the parameters of the Wilson–Cowan model and the numerical values used for the simulations. For parameters that were varied across simulations, the value reported here corresponds to the value used in the main simulation setting, and any deviations from this value are explicitly stated in the main text.

| Parameter | Description | Value |
|---|---|---|
| $\tau_E$ | Excitatory population time constant | 2.5 ms |
| $\tau_I$ | Inhibitory population time constant | 3.75 ms |
| $c_{EE}$ | Local excitatory-to-excitatory coupling | 16 |
| $c_{EI}$ | Local excitatory-to-inhibitory coupling | 15 |
| $c_{IE}$ | Local inhibitory-to-excitatory coupling | 12 |
| $c_{II}$ | Local inhibitory-to-inhibitory coupling | 3 |
| $a_E$ | Gain of excitatory sigmoid | 1.5 |
| $a_I$ | Gain of inhibitory sigmoid | 1.5 |
| $\mu_E$ | Midpoint of the excitatory sigmoid | 3 |
| $\mu_I$ | Midpoint of the inhibitory sigmoid | 3 |
| $P_E$ | Baseline external input to the excitatory population | 0.55 |
| $P_I$ | Baseline external input to the inhibitory population | 0 |
| $\tau_{OU}$ | Correlation timescale of the Ornstein-Uhlenbeck process | 5 ms |
| $\sigma_{OU}$ | Standard deviation of the Ornstein-Uhlenbeck process | $10^{-4}$ |
| $\mu_{OU}$ | Mean of the Ornstein-Uhlenbeck process | 0 |
| $s$ | Common-driver strength: fraction of the stochastic OU drive shared across all nodes | 0; **varied** |
| $G$ | Global coupling scaling factor | 0.1; **varied** |
| $dt$ | Integration time step | 0.1 ms |

**Table S2 Parameters of the Wilson-Cowan model with spike-frequency adaptation.** The table lists the parameters used in The Virtual Brain (TVB) implementation of the WCA model. All parameter values were taken from (2), except for the thresholds of the inhibitory input-output function ($h_i$) and of adaptation ($\mu_a$), which were decreased from 12 to 5 and from 1.5 to 0.5, respectively, to increase the contribution of the inhibitory and adaptation processes.

| Parameter | Description | Value |
|---|---|---|
| $\tau_{e/i}$ | Time constant of the excitatory/inhibitory population | 5 ms |
| $\tau_a$ | Time constant of the adaptation dynamics | 500 ms |
| $w_{ee}$ | Local excitatory-to-excitatory coupling | 4 |
| $w_{ei}$ | Local excitatory-to-inhibitory coupling | 4 |
| $w_{ie}$ | Local inhibitory-to-excitatory coupling | 3 |
| $w_{ii}$ | Local inhibitory-to-inhibitory coupling | 0 |
| $k_{e/i}$ | Amplitude or scaling factor for the excitatory/inhibitory input-output function | 0.02/0.05 |
| $h_{e/i}$ | Threshold of excitatory/inhibitory input-output function | 0/5 |
| $n$ | Power law exponent of input-output functions; determines the sharpness of the population response | 2 |
| $a_a$ | Gain of the adaptation dynamics | 5 |
| $\mu_a$ | Firing threshold of the adaptation | 0.5 |
| $g$ | Adaptation strength | 1.8; 3.6 |
| $I_{e/i}$ | External input to excitatory/inhibitory population | 3.6/3.6 |
| $\tau_{OU}$ | Correlation timescale of the Ornstein-Uhlenbeck process | 5 ms |
| $\sigma_{OU}$ | Standard deviation of the Ornstein-Uhlenbeck process | 0.001 |
| $\mu_{e/i}$ | Mean of the Ornstein-Uhlenbeck process | 0/0 |
| $c$ | Factor scaling the excitatory coupling strength between nodes | 0.15 |
| $dt$ | Integration time step | 0.5 ms |

### 4. Supplementary Notes

##### 4.1 Perturbational complexity reflects small-world organization beyond local attractor stability

As shown in Fig. 1D (main results), clustered motifs altered the attractor landscape across network architectures. Clustered and balanced architectures, both characterized by high clustering, exhibited similar relaxation times $\langle\tau\rangle$, whereas random architectures showed shorter $\langle\tau\rangle$. Combined with the higher perturbational complexity of balanced relative to clustered networks (Fig. 3), this indicates that differences in proximity to the stability boundary cannot account for the advantage of balanced architectures over clustered ones. By contrast, the lower perturbational complexity of random networks could still partly reflect their greater distance from the stability boundary and the resulting faster decay of evoked reverberations (Supplementary Fig. S10). We therefore examined whether the association between structural complexity and perturbational complexity could be indirect and largely explained by differences in the local stability of individual nodes.
We first exploited the time-resolved formulation of PCI (12) to quantify complexity before the faster decay of random networks could substantially penalize them. PCI was evaluated in the first 100 ms after stimulation (Fig. S11A). Even at this early stage, PCI remained higher in balanced than in random architectures and was robustly associated with the small-world propensity of the networks (randomization: $\rho=0.81$; modularization: $\rho=0.78$; Fig. S11B).
This observation motivated a relaxation time-matched analysis (Fig. S11C) to further control for differences in proximity to the stability boundary. We retained only networks whose mean relaxation time, $\langle\tau\rangle$, did not exceed the maximum value observed among the random networks R ($R\langle\tau\rangle$,max=41.4). This yielded a subset comprising the 10 R networks and 71 additional networks from the randomization and modularization families. In this subset, the relaxation time distributions of the non-random networks (mean ± SD = 39.7 ± 1.1) and R networks (39.3 ± 0.9) did not differ (Kruskal–Wallis test, p = 0.21). Likewise, the mean late evoked activity ($mean_{LA}$) was comparable across groups (non-random: 9.9 ± 4.4; R: 8.4 ± 3.4; Kruskal–Wallis test, p = 0.27). Remarkably, after matching relaxation dynamics, observational signal diversity measures remained unrelated to small-world propensity, whereas interaction-based measures (NC and $F_{comp}$) showed a positive association with it (Fig. S11D). A similarly robust positive relationship was observed for all perturbational complexity measures.

Together, these findings indicate that differences in local attractor stability may constitute one mechanism through which topology influences perturbational complexity, but they do not fully account for the observed effects (see Supplementary Notes 4.2). After controlling for effective proximity to the stability boundary, small-world organization remained associated with higher perturbational complexity as well as higher observational complexity based on network interaction.

##### 4.2 Local perturbation gain, network amplification, and attractor stability

A potential concern when comparing evoked responses across network topologies is that the same external perturbation may produce different effective displacements from the fixed point if the networks differ in their stability properties. Here, it is important to distinguish the local susceptibility to the external input from the network-level amplification generated by recurrent connectivity.

Across all network architectures, local node parameters, the operating fixed point, and the slope of the transfer function at that fixed point were held constant. Therefore, for a given external input, the directly stimulated node has the same local input–output gain across topologies. Linearizing the dynamics around the common fixed point $x*$ gives

$$\delta\dot{x} = J_G \delta x + B\,u(t),$$

where $u(t)$ denotes the external perturbation, $B$ its direct coupling to the stimulated node, and $J_G$ is the Jacobian of the full network, which depends on topology $G$. Because the local operating point and node dynamics are fixed, differences in $J_G$ arise from the organization of recurrent interactions rather than from differences in local excitability.

Importantly, equal local gain does not imply an identical network-level response. For a finite perturbation,

$$\delta x(t) = \int_0^t e^{J_G(t-s)} B\,u(s)\,ds,$$

so the same external input can generate different distributed responses because the network's linear evolution operator $e^{J_G t}$ depends on topology. Differences in network-level response magnitude, propagation, and reverberation therefore do not reflect differences in the applied input, but are part of the causal effect of topology itself: connectivity determines how the initially injected activity is redistributed, amplified, and recirculated through recurrent pathways.

Topology also altered effective proximity to the stability boundary, as reflected by differences in the relaxation time $\tau$. This introduces a related but distinct effect: networks with slower relaxation can sustain evoked activity for longer, potentially increasing perturbational complexity simply by extending the temporal window available for recurrent interactions. The analyses in Fig. S11 were designed to separate this contribution from the broader effect of topology. First, the time-resolved PCI analysis showed that the advantage of balanced architectures was already present within the first 100 ms after stimulation, before differences in late response decay could fully account for it. Second, restricting the analysis to networks with comparable ($\langle\tau\rangle$) controlled for differences in effective proximity to the stability boundary as indexed by the dominant relaxation timescale. Within this matched subset, perturbational complexity remained positively associated with small-world propensity.

Matching ($\tau$) does not, and is not intended to, make the complete network transfer function identical across architectures. Networks with comparable dominant relaxation times may still differ in their eigenspectra, eigenvectors, recurrent pathways, and transient amplification. These residual differences are precisely among the mechanisms through which topology can shape the response to an otherwise locally equivalent perturbation. Thus, the present controls isolate two potential trivial explanations (i.e., different local input-output gain and different dominant relaxation dynamics) while preserving the topology-dependent network interactions whose contribution to perturbational complexity is the object of study.
